# Reduced antibacterial drug resistance and *bla*_CTX-M_ β-lactamase gene carriage in cattle-associated *Escherichia coli* at low temperatures, at sites dominated by older animals and on pastureland: implications for surveillance

**DOI:** 10.1101/778407

**Authors:** Hannah Schubert, Katy Morley, Emma F. Puddy, Robert Arbon, Jacqueline Findlay, Oliver Mounsey, Virginia C. Gould, Lucy Vass, Madeleine Evans, Gwen M. Rees, David C. Barrett, Katy M. Turner, Tristan A. Cogan, Matthew B. Avison, Kristen K. Reyher

## Abstract

Little is known about the drivers of critically important antibacterial resistance in species with zoonotic potential present on farms (e.g. CTX-M □-lactamase-positive *Escherichia coli*). Here, we collected samples, monthly over a two-year period, on 53 dairy farms in the South West of England, and data for 610 variables concerning antimicrobial usage, management practices and meteorological factors. We detected *E. coli* resistant to amoxicillin, ciprofloxacin streptomycin and tetracycline, respectively, in 2754/4145 (66%), 263/4145 (6%), 1475/4145 (36%) and 2874/4145 (69%) of all samples from faecally contaminated sites. *E. coli* positive for *bla*_CTX-M_ were detected in 224/4145 (5.4%) of samples. Multilevel, multivariable logistic regression showed antibiotic dry cow therapeutic choice (including use of cefquinome or framycetin) to be associated with increased odds of *bla*_CTX-M_ positivity. Low temperature was associated with reduced odds of *bla*_CTX-M_ *E. coli* positivity in samples and to reduced odds of finding *E. coli* resistant to each of the four test antibacterials. This was additional to the effect of temperature on total *E. coli* density. Furthermore, samples collected close to calves had increased odds of having *E. coli* resistance to each antibacterial or positive for *bla*_CTX-M_. Samples collected on pastureland had reduced odds of having *E. coli* resistant to amoxicillin or tetracycline, and being positive for *bla*_CTX-M_.

**Importance:** Antibacterial resistance poses a significant threat to human and animal health and global food security. Surveillance for resistance on farms is important for many reasons, including to track the impacts of interventions aimed at reducing the prevalence of resistance. In this epidemiological survey of dairy farm antibacterial resistance, we show that local temperature, as it changes over the course of a year, is associated with the prevalence of antibacterial resistant *E. coli*. Also, that prevalence of resistant *E. coli* is higher in indoor environments and in environments inhabited by young animals. These findings have profound implications for routine surveillance and for surveys carried out for research. They provide important evidence that sampling at a single time-point and/or single location on a farm is unlikely to be adequate to accurately determine the status of the farm with regard to the presence or number of resistant *E. coli*.

## Introduction

Antimicrobial resistance - and particularly antibacterial resistance (ABR) - is a significant global challenge. Many countries are implementing plans to reduce the use of antibacterial drugs (ABs) in food-producing animals. For example, the most recent UK five-year National Action Plan includes a target to reduce AB use (ABU) in the treatment of food-producing animals by 25% (1). In Europe, AB sales for food-producing animals fell by 20% from 2011 to 2016 (2). In the UK dairy industry, overall ABU dropped from 24 mg/kg in 2015 to 17 mg/kg in 2018 (3, 4). In 2018, additional industry-led policies were enforced in the UK that aimed to almost eliminate the use of highest-priority critically important antimicrobials (HP-CIAs) such as third- and fourth-generation cephalosporins (3GCs and 4GCs) as well as fluoroquinolones on dairy farms. One reason for reducing ABU in farming is to reduce the prevalence of ABR bacteria carried by farm animals. However, there is a need for better data on drivers of ABR in farming. More granularity of understanding is required concerning the risks of using individual ABs and other management practices. This is especially important in terms of drivers of HP-CIA resistance. A focus among HP-CIAs is on 3GC and fluoroquinolone resistance in *Escherichia coli*, a species commonly found in animal faeces and considered one of the most significant potential zoonotic threats to humans (5).

3GC resistance is increasingly prevalent in *E. coli* causing infections in humans (6) and is also found in farmed and domestic animals around the world (7). The production of CTX-M (an extended-spectrum □-lactamase) is the most common mechanism of 3GC resistance in *E. coli* in humans in the UK; for example, in a recent study of urinary *E. coli* from humans in South West England, 82.2% of 3GC-resistant isolates carried *bla*_CTX-M_ (8).

The objective of this study was to describe the prevalence of 3GC-resistant *E. coli* carrying *bla*_CTX-M_ as well as *E. coli* resistant to amoxicillin, tetracycline, streptomycin and the fluoroquinolone ciprofloxacin found in faecally contaminated environments of dairy cattle in a geographically restricted population of dairy farms in South West England. Furthermore, this study investigated environmental, ABU and management practice risk factors for the presence of such *E. coli.*

## Results

### Prevalence and PCR characterisation of 3GC-resistant *E. coli* from dairy farms

4581 samples from faecally contaminated sites were collected from 53 dairy farms. Samples were collected on each farm monthly between January 2017 and December 2018. 4145 samples were positive for growth of *E. coli* on non-selective agar. Of these, 384/4145 (9.3%) samples representing 47/53 (88.7%) of farms were positive for growth of *E. coli* on agar containing the 3GC cefotaxime. From these, 1226 3GC-resistant isolates were taken forward for PCR testing for possible cephalosporinase genes of interest (GOIs): *bla*_CTX-M_ (groups 1, 2, 8, 9 and 25), *bla*_CMY_, *bla*_DHA_ and *bla*_SHV_. Over half (648/1226; 52.7%) of all isolates tested were found to harbour *bla*_CTX-M_ genes. Of these, 547/648 (84.4%) were of group 1, 99/648 (15.3%) were of group 9, and, in one case, both gene groups were identified. Twelve isolates harboured a *bla*_CMY_ gene – one alongside *bla*_CTX-M_ group 1 – and one isolate was *bla*_DHA-1_-positive. No isolates were positive for *bla*_SHV_ and the remaining 566/1226 (46.2%) isolates were PCR-negative for all GOIs. These isolates were hypothesised to hyper-produce the chromosomally encoded AmpC β-lactamase; some of these isolates have been characterised in detail in a separate study (9).

### Farm- and sample-level risk factors for *bla*_CTX-M_ *E. coli* positivity

Based on PCR, carriage of *bla*_CTX-M_ was the most common mechanism of 3GC-resistance in *E. coli* from dairy farms in this study. Identifying management practice- and AMU-associated risk factors for *bla*_CTX-M_ *E. coli* positivity was therefore considered to be an important objective. Overall, 5.4% (224/4145) of samples representing 42/53 (79.2%) of farms contained 3GC-resistant *E. coli* confirmed to carry *bla*_CTX-M_ using PCR. Positivity for *bla*_CTX-M_ *E. coli* was three times higher in Calf samples (samples collected from the environments of calves) (98/631 [15.5%] of samples) than overall (**Table 1**).

**Table 1.** Prevalence of *E. coli* carrying *bla*_CTX-M_ at farm and sample levels

| Sample type |  | Farm level | Sample level |
| --- | --- | --- | --- |
| Overall | Total sample size | 53 | 4145 |
|  | Total (%) positive for CTX-M-carrying *E. coli* | 42 (79.2%) | 224 (5.4%) |
| Adult | Total sample size | 52 | 1835 |
|  | Total (%) positive for CTX-M-carrying *E. coli* | 25 (48.1%) | 76 (4.1%) |
| Dry Cow | Total sample size | 46 | 282 |
|  | Total (%) positive for CTX-M-carrying *E. coli* | 7 (15.2%) | 8 (2.8%) |
| Calf | Total sample size | 51 | 631 |
|  | Total (%) positive for CTX-M-carrying *E. coli* | 33 (64.7%) | 98 (15.5%) |
| Heifer | Total sample size | 41 | 1235 |
|  | Total (%) positive for CTX-M-carrying *E. coli* | 18 (44%) | 40 (3.2%) |
| Pastureland | Total sample size | 47 | 630 |
|  | Total (%) positive for CTX-M-carrying *E. coli* | 8 (17%) | 12 (1.9%) |
| Pastureland that is publicly accessible (Footpath) | Total sample size | 41 | 395 |
|  | Total (%) positive for CTX-M-carrying *E. coli* | 8 (20.0%) | 11 (2.8%) |

Given the high positivity rate for *bla*_CTX-M_ *E. coli* in Calf samples, a separate risk factor analysis using only Calf data was performed. One farm-level fixed effect and three sample-level fixed effects were retained in the final multilevel, multivariable logistic regression model (**Table S1, Table 2**). The use of cefquinome or framycetin dry cow therapies were both associated with increased odds of *bla*_CTX-M_ *E. coli* positivity, as was higher average monthly temperature. Plotting sample-level positivity for *E. coli* carrying *bla*_CTX-M_ versus average monthly temperature revealed that the relationship between positivity and temperature was primarily driven by low *bla*_CTX-M_ *E. coli* positivity rates in months where the average temperature was below 10°C (**Figure 1A**).

**Table 2.** Fixed effects from the multilevel, multivariable logistic regression model predicting *bla*_CTX-M_ *E. coli* positivity performed on Calf samples

| Risk factor | Odds ratio [95% confidence interval] | p |
| --- | --- | --- |
| Use of cefquinome dry cow therapy in the last six months | 4.18 [2.11, 8.25] | 0.00003 |
| Daily water trough cleaning | 0.44 [0.29, 0.69] | 0.0002 |
| Average monthly temperature | 1.57 [1.20, 2.06] | 0.0008 |
| Use of framycetin dry cow therapy in the last six months | 1.91 [1.01, 3.61] | 0.04 |

**Figure 1.**
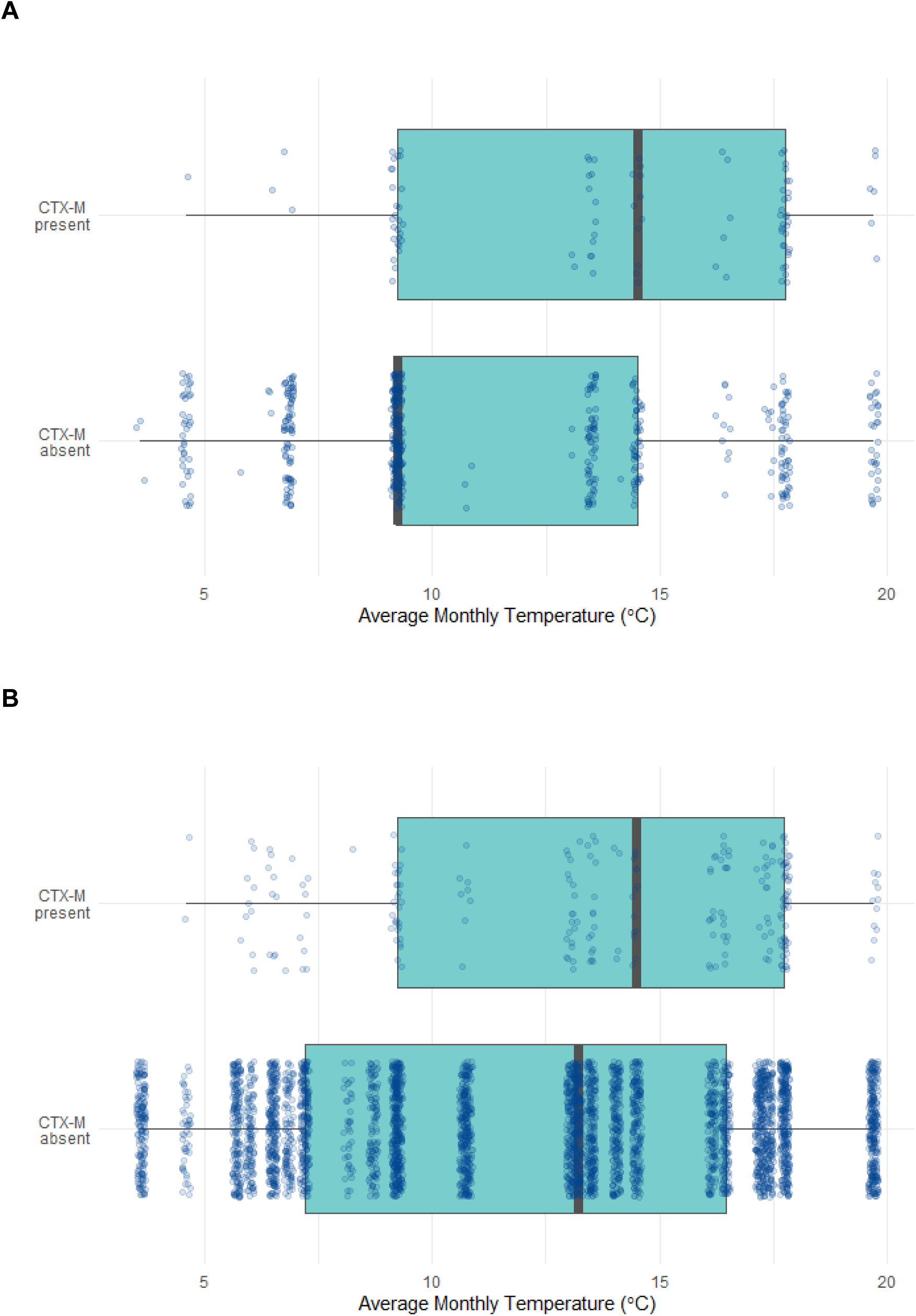
Average monthly temperature vs. presence (1) or absence (0) of *E. coli* positive for *bla*_CTX-M_ in samples from (**A)** pre-weaned calves and (**B**) all faecally contaminated dairy farm environments. Each sample is represented by a dot. A multilevel, multivariable logistic regression model revealed a positive association with increased temperature in both cases (p=<0.0001).

Risk factor analysis was also performed for the full dataset. One farm-level fixed effect and three sample-level fixed effects were retained in the final model (**Table S2, Table 3**). Interestingly, this model revealed that *bla*_CTX-M_ *E. coli* was less likely to be found in samples obtained from pastureland, which included publicly accessible farmland (Footpaths) compared with other sample types. Analysis of the full dataset confirmed what was seen with the Calf dataset: higher average monthly temperature was associated with an increased odds of *bla*_CTX-M_ *E. coli* positivity. Again, visualisation of the data confirmed that this was primarily driven by a reduction in *bla*_CTX-M_ *E. coli* positivity rate in months with an average temperature below 10°C (**Figure 1B**).

**Table 3.** Fixed effects from the multilevel, multivariable logistic regression model predicting *bla*_CTX-M_ *E. coli* positivity performed on the full dataset

| Risk factor | Odds ratio [95% confidence interval] | p |
| --- | --- | --- |
| Sample taken from the environment of pre-weaned heifers | 4.52 [3.25, 6.27] | <0.00000001 |
| Average monthly temperature | 1.61 [1.36, 1.90] | 0.00000001 |
| Sample taken from pastureland | 0.32 [0.17,0.61] | 0.0004 |
| Feeding of maize silage | 3.28 [1.50, 7.18] | 0.002 |

A Bayesian logistic regression model was also constructed in which the effect of total farm ABU and specifically total 3GC and 4GC use were tested as predictors for *bla*_CTX-M_ *E. coli* positivity in the total dataset, with 102 potential confounders included. The impact of temperature (odds ratio 1.71 [1.42, 2.05]) on *bla*_CTX-M_ *E. coli* positivity was also retained in this alternative model (**Table S3**).

Defining sample-level positivity for *bla*_CTX-M_ *E. coli* is dependent upon finding *bla*_CTX-M_ using PCR in *E. coli* colonies that have grown on agar containing cefotaxime. If *bla*_CTX-M_ *E. coli* in a sample exist at such a low density that they are not detected using selective agar, the sample will be falsely identified as negative for *bla*_CTX-M_ *E. coli*. This impact of bacterial density on assay sensitivity is an important consideration in the context of the finding that *bla*_CTX-M_ positivity is low at low temperatures. To account for this, the logistic link function was adjusted (see Supplementary). This only modestly altered the effect sizes and the p-values for the risk factors (**Figure S2**), confirming that the effect of low temperature on *bla*_CTX-M_ *E. coli* positivity was additional to its effect on *E. coli* prevalence. All values in **Tables 2** and **3** come from models with this adjusted logistic link function applied.

### Prevalence and risk factor analysis for *E. coli* resistant to other antibacterial classes

All 4145 samples positive for growth of *E. coli* on non-selective agar were also tested for detectable numbers of *E. coli* resistant to four non-cephalosporins: amoxicillin, tetracycline, streptomycin and ciprofloxacin, the last being representative of the HP-CIA class, the fluoroquinolones. Resistance to amoxicillin and tetracycline were the most prevalent types of resistance found, with ciprofloxacin resistance being the least commonly detected (**Table 4**).

**Table 4.** Farm and sample-level prevalence of resistance to non-cephalosporins

| Antibacterial drug | Farm level resistance | Sample level resistance |
| --- | --- | --- |
| Amoxicillin | 53/53 (100%) | 2754/4145 (66%) |
| Ciprofloxacin | 49/53 (92%) | 263/4145 (6%) |
| Streptomycin | 53/53 (100%) | 1475/4145 (36%) |
| Tetracycline | 53/53 (100%) | 2874/4145 (69%) |

Using a Bayesian logistic regression method, factors associated with the risk of a sample being positive for *E. coli* resistant to each of the test ABs were identified from the total dataset. As seen for *bla*_CTX-M_ *E. coli*, where and when the samples were collected were more significantly associated with the odds of finding resistant *E. coli* in a sample than farm-level management practices or ABU, with all four models showing a positive association between average monthly temperature and the odds of finding resistant *E. coli* in a sample (**Table 5**). Also consistent across all models was the significance of sampling different areas of the farm. Again, as with *bla*_CTX-M_ *E. coli*, environments of calves were more likely (as a proportion of all *E. coli* than elsewhere on the farm) to harbour all four types of resistant bacteria. Samples collected from pastureland were, like *bla*_CTX-M_ *E. coli*, negatively associated with the presence of amoxicillin and tetracycline resistance (**Table 5**).

**Table 5.** Fixed effects from a Bayesian model predicting resistant *E. coli* positivity performed on the full dataset

| Risk factor | Odds ratio [95% credible interval] |
| --- | --- |
| **Amoxicillin resistance** |  |
| Average monthly temperature | 1.91 [1.57, 2.35] |
| Sample taken from the environment of pre-weaned heifers | 1.99 [1.29, 2.98] |
| Sample taken from pastureland | 0.27 [0.20, 0.37] |
| **Ciprofloxacin resistance** |  |
| Average monthly temperature | 2.14 [1.63, 2.87] |
| Sample taken from the environment of pre-weaned heifers | 4.13 [2.79, 6.46] |
| **Streptomycin resistance** |  |
| Average monthly temperature | 1.53 [1.32, 1.77] |
| Sample taken from the environment of pre-weaned heifers | 1.95 [1.46, 2.51] |
| **Tetracycline resistance** |  |
| Average monthly temperature | 1.98 [1.55, 2.55] |
| Sample taken from the environment of pre-weaned heifers | 3.39 [2.00, 5.82] |
| Sample taken from pastureland | 0.24 [0.15, 0.35] |
| Sample taken during the calving season | 2.02 [1.10, 3.29] |
| Average monthly rainfall | 1.26 [1.08, 1.47] |
| Total farm streptomycin use in 2017 measured in mg/PCU* | 0.76 [0.60, 0.97] |
| Total farm ABU in 2017 measured in mg/PCU* | 1.47 [1.01, 2.13] |

Full results for all variables tested can be found in **Table S4**. Re-running the models with sceptical priors did not affect the results; only very small differences in the model coefficients were observed (**Table S5**). Details of all model checking can be found in Supplementary.

## Discussion

### Prevalence of *bla*_CTX-M_ positive *E. coli*

This study is unique in its scale: extensive management practice and ABU data along with multiple samples from multiple farms were collected monthly across a two-year period. Overall, 224/4145 (5.4%) of samples were positive for *E. coli* carrying *bla*_CTX-M_. This compares well with previously calculated *bla*_CTX-M_ *E. coli* carriage of approximately 7% in Danish slaughter pigs (10) and 3.6% in UK broiler chickens and turkeys (11). Various studies have identified much higher prevalence in chicken meat, but this could be due to cross-contamination at slaughter and in the food chain (12, 13).

Studies examining the prevalence of *bla*_CTX-M_ *E. coli* in human populations have shown mixed results. A prevalence of 65.7% was found amongst commensal isolates in Thailand (14). In the UK, a study across four regions reported commensal faecal carriage of *bla*_CTX-M_ *E. coli* to be approximately 7% (15). A recent analysis of human urinary samples from the same region as the farms surveyed in this study gave a sample-level prevalence of *bla*_CTX-M_ *E. coli* of approximately 5% (8). It should be noted that all farm samples in the present study were from faecally contaminated sites, not individual animals, and so it is possible that the number of animals carrying *bla*_CTX-M_ *E. coli* was much lower than the reported sample-level prevalence. Direct comparison with human and other farm animal carriage studies should therefore be made with caution.

### Impact of temperature on the odds of finding resistant and *bla*_CTX-M_-positive *E. coli*

This study found 42/53 (79%) of farms to be positive, based on phenotypic analysis and PCR, for *bla*_CTX-M_ *E. coli*. This was higher than seen in other using similar methodology e.g. 17/48 (35%) of randomly selected UK dairy farms (16) and 5/25 (20%) of farms in Ohio (17) were positive. In the present study, samples were collected each month over two years, hence the chances of finding a positive sample on each farm may have been greater than in these earlier point-prevalence studies (16,17). When farm-level positivity for *bla*_CTX-M_ *E. coli* was plotted on a month-by-month basis (**Figure 2**) the highest prevalence for a single monthly survey was 22.5%, which fits more closely with these other studies.

**Figure 2.**
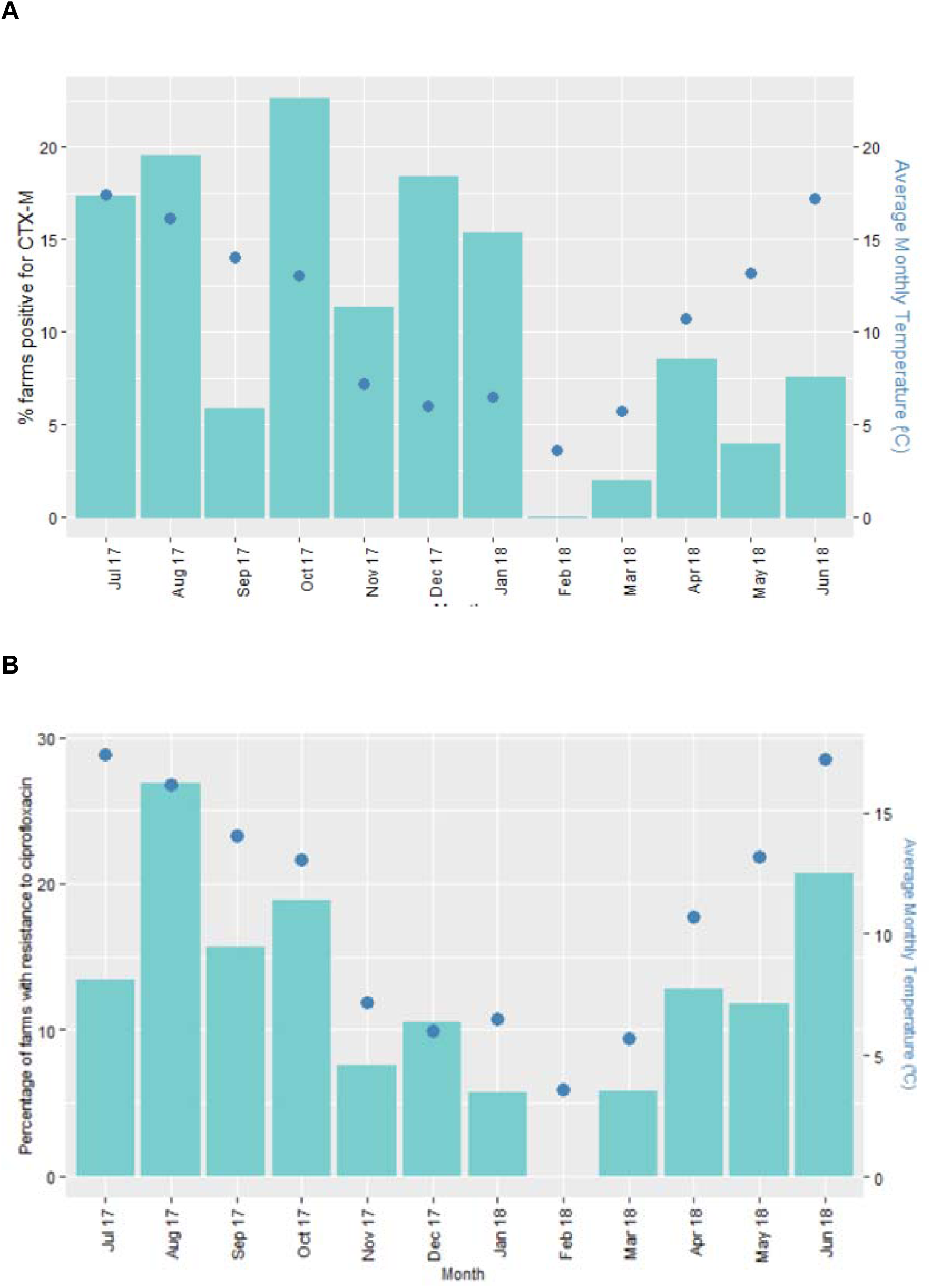
Percentage of farms positive for (**A**) *E. coli* positive for *bla*_CTX-M_ in samples (**B**) ciprofloxacin resistant *E. coli*. Data are presented by month (bars) and overlayed by a graph of average monthly temperature (dots) representing a year during the middle period of this study. Samples from calves have been excluded.

Sample-level prevalence of *bla*_CTX-M_ *E. coli* was low overall (5.4%). This contrasts with >90% (18) or 50% (19) of *bla*_CTX-M_ *E. coli* in samples taken from bovine faecal pats. This difference could be due to the large number of samples collected here, particularly over winter, given low temperature was associated with low *bla*_CTX-M_ *E. coli* positivity (**Fig. 1A; 1B**). Indeed, the observation that average monthly temperature had a significant effect on *bla*_CTX-M_ *E. coli* positivity (**Table 3**), as well as positivity for resistance to amoxicillin, ciprofloxacin, streptomycin and tetracycline (**Table 5**) highlights problems with studies where a single time-point or sampling season is used. **Figure 2** shows the stark impact of this in real terms: *bla*_CTX-M_ *E. coli* positivity and positivity for ciprofloxacin-resistant *E. coli* at farm level was zero in February, the coldest month of the year (based on average temperature; **Fig 2A, B**). Whilst average annual temperature found at locations across an entire continent has previously been shown to impact average ABR levels at those locations (20), the finding that periods of low temperatures were associated with lower prevalence of a dominant cause of ABR - and particularly HP-CIA resistance at a given location during the course of a year - is particularly important. This observation also leads to concern about the impact of climate change - and especially increasing temperatures - on attempts to reduce ABR. Whilst temperature was associated with the total number of *E. coli* found in each sample, this was accounted for using a measurement error method incorporated into the model; as such, the effect of temperature on ABR or *bla*_CTX-M_-positive *E. coli*, whilst in part driven by the effect on total *E. coli* number, also had an independent association suggestive of a temperature-dependent fitness burden of resistance.

### High levels of ABR and *bla*_CTX-M_ positive *E. coli* in farm locations dominated by young animals and low levels on pastureland

There were clear differences in the risk of encountering *bla*_CTX-M_ *E. coli* at different sites on a farm (e.g. 15.5% in Calf samples, 4.1% in Adult samples). The Calf environment was also much more likely to have *E. coli* resistant to amoxicillin, tetracycline, streptomycin, and ciprofloxacin, so this seems to be a universal effect. Other studies have also generally found high levels of resistance in samples collected from or in the environment of younger calves (21-24). There may also be an association with temperature here since calves are generally kept in warmer environments than are adult cows but may also be due to some physiological change with age.

This study also identified a decreased odds of detecting *bla*_CTX-M_ *E. coli* in samples collected on pastureland compared with those collected elsewhere on the farm. This relationship also held for *E. coli* resistant to amoxicillin and tetracycline. Because pastureland may be more affected by the elements, this finding may be partly linked with the association between temperature and ABR.

### AB contamination of colostrum as a possible driver of *bla*_CTX-M_ *E. coli* positivity in dairy calves - evidence of direct and co-selection

It has been shown experimentally that feeding waste (AB-contaminated) milk to calves increases faecal excretion of ABR bacteria (25). This practice is reducing on UK dairy farms and, in the analysis presented here, waste milk feeding was not associated with an increased risk of finding ABR or *bla*_CTX-M_ positive *E. coli*. In contrast, the choice of dry cow therapy (an AB preparation inserted into a cow’s udder between lactations to help treat or prevent mastitis) was associated with *bla*_CTX-M_ *E. coli* positivity in Calf samples (**Table 2**). It has previously been shown that colostrum from cows given cefquinome dry cow therapy is heavily contaminated with cefquinome (26), and colostrum management is a hugely important part of early life for most farmed mammals and is universally encouraged in dairy farming. In this study, cefquinome (a 4GC) dry cow therapy was most significantly associated with *bla*_CTX-M_ *E. coli* in Calf samples (**Table 2**). This can be explained by direct selection because production of CTX-M confers 4GC resistance in *E. coli* (27). There was also a clear positive association between the usage of framycetin as part of a dry cow therapy combination and the odds of finding *bla*_CTX-M_ *E. coli* in Calf samples (**Table 2**). Whilst no work has been published on the contamination of colostrum with framycetin, its use as a mastitis therapy for milking cows results in identifiable residues in milk (28), so it is highly likely to also contaminate colostrum. It is possible, therefore, that feeding of colostrum - which can be contaminated with AB used for dry cow therapy - is a driver of *bla*_CTX-M_ *E. coli* in calves. An alternative (or indeed an additional) explanation for this observed association is that *E. coli* (a species known to be found in the udders of dairy cows; 29) that carry *bla*_CTX-M_ are selected within the udder during AB dry cow therapy and contaminate colostrum alongside the AB used. Others (16) also identified overall use of 3/4GCs as a risk factor for *bla*_CTX-M_ *E. coli* presence on dairy farms but have not made a link between the usage of framycetin and prevalence of *bla*_CTX-M_ *E. coli*. However, it is not always clear whether other studies have separated out different dry cow therapies since they have tended to focus on systemic AB use. Clearly, an aminoglycoside like framycetin cannot directly select for *bla*_CTX-M_ *E. coli*, but aminoglycoside resistance genes are common on plasmids (30), as is *bla*_CTX-M_ (27) and so this may be an example of co-selection, where selection of resistance to one antibacterial class increases resistance to another.

## Conclusions

Overall, we provide important evidence that sampling at single time-points and/or limited locations on a farm are unlikely to be adequate to accurately determine the status of the farm concerning the prevalence of ABR *E. coli*. This makes comparisons between surveillance studies on farms, designed for research or for regulatory purposes, extremely difficult, and we urge for a standardised, multi-sample framework accounting for the differential risk factors identified here to be used in the design of future studies.

## Materials and Methods

### Farm recruitment and ethical approval

A convenience sample of 53 dairy farms was recruited through personal contacts, local veterinary practices, and milk processors. Details of the study population are presented in Supplementary. Of these, 43 farms were in a 50 x 50 km area defined based on the locations of 146 general practices that referred routine urine samples from human patients to the microbiology reference lab at Severn Pathology, Southmead Hospital (8). A further 10 study farms were clustered in a separate region in South West England. All farmers gave fully informed consent to participate in the study. Ethical approval was obtained from the University of Bristol’s Faculty of Health Sciences Research Ethics Committee (ref 41562).

### Farm sampling and sample processing

Farms were visited monthly between January 2017 and December 2018. Samples were collected using sterile overshoes (over-boot socks) traversing farm areas. Where access was restricted (e.g. for pens containing single or pairs of calves), samples were collected directly from the ground using gloved hands. Details of the six types of samples collected are in Supplementary; these represent faecally contaminated environments representative of milking cows (Adult), cows between periods of lactation (Dry Cow), heifers after weaning (Heifer), heifers before weaning (Calf). Samples of these types collected from pastureland were designated as such for separate analysis (Pasture) which also included samples collected from publicly accessible land on the farm (Footpath). Samples were refrigerated from collection to processing, were transferred into individual labelled sterile stomacher bags, and suspended in 10 ml.g^-1^ of phosphate buffered saline (PBS Dulbecco A; Oxoid, Basingstoke, UK). Samples were then mixed for one min in a stomacher (Stomacher 400, Seward, Worthing, UK). Samples were mixed 50:50 with sterile glycerol and aliquots stored at -80°C.

### Microbiology and PCR analysis

Twenty microlitres of sample (diluted 1:10) were spread onto tryptone bile X-glucuronide agar (TBX; Scientific Laboratory Supplies); 20 µL of undiluted sample were spread onto TBX agar containing 16 mg.L^-1^ tetracycline, 8 mg.L^-1^ amoxicillin, 0.5 mg.L^-1^ ciprofloxacin, 16 mg.L^-1^ streptomycin or 16 mg.L^-1^ cephalexin. Plates were incubated at 37°C, and the number of blue colonies (*E. coli*) counted. Samples yielding no *E. coli* colonies on antibiotic-free agar were excluded from further analysis. Up to five *E. coli* isolates from each cephalexin TBX agar plate were transferred onto cefotaxime (CTX, 2 mg.L^-1^) TBX agar. All AB concentrations were chosen as those which define clinically relevant resistance in humans according to EUCAST (31). Two multiplex PCRs were performed to screen for β-lactamase genes in CTX-R *E. coli*. The first was to detect *bla*_CTX-M_ groups as previously described (32) and the second was to detect the following additional β-lactamase genes: *bla*_CMY_, *bla*_DHA_, *bla*_SHV_, *bla*_TEM_, *bla*_OXA-1_ (8).

### Risk factor analyses

The risk factors examined fell into four categories: farm management, ABU, sample characteristics and meteorological. Four management practice questionnaires were developed (details provided in Supplementary). ABU was extracted from prescribing and sales data between Jan 2016 to Dec 2018 obtained from veterinary practices servicing the study farms. For two farms, sales data were not available and on-farm records were used. The full method for obtaining, cleaning, and processing ABU data can be found in Supplementary. Local meteorological data were extracted from publicly available UK Met Office data (https://www.metoffice.gov.uk/pub/data/weather/uk/climate/stationdata/yeoviltondata.txt).

Sample processing and data analysis workflows are illustrated in **Figure S1**. All data analysis was performed using R (https://www.r-project.org/). Two modelling approaches were used: 1) variable selection via univariable screening and stepwise model selection with a multilevel, multivariable logistic regression model and 2) a regularised Bayesian model. Both were used to analyse risk factors associated with *bla*_CTX-M_ *E. coli* positivity; the second was also used to analyse risk factors associated with positivity for *E. coli* resistant to amoxicillin, ciprofloxacin, streptomycin, and tetracycline. Sensitivity analyses were performed to test for measurement bias, to account for the fact that resistant *E. coli* were more likely to be found in a sample if there was a higher density of bacterial colony forming units. Further details of variable selection and development of the models and model checking are provided in Supplementary.

## Acknowledgements

We wish to thank all the farmers and veterinary surgeons who participated in this study. In addition, we would like to thank the milk processing companies who supported our recruitment process and the veterinary students who helped with data processing and sample collection. Genome sequencing was provided by MicrobesNG (http://www.microbesng.uk), which is supported by the BBSRC (grant number BB/L024209/1).

## Funding

This work was funded by grant NE/N01961X/1 to M.B.A., K.K.R., K.M.T., T.A.C. and D.C.B. from the Antimicrobial Resistance Cross Council Initiative supported by the seven United Kingdom research councils. R.A. was supported by the Jean Golding Institute for data-intensive research at the University of Bristol. G.M.R. was funded by the Langford Trust.

## Transparency declarations

D.C.B. was president of the British Cattle Veterinary Association 2018-19. Otherwise, the authors declare no competing interests. Farming and veterinary businesses who contributed data and permitted access for sample collection were not involved in the design of this study or in data analysis and were not involved in drafting the manuscript for publication.

## Author Contributions

Conceived the Study: D.C.B., K.K.R., M.B.A.

Collection of Data: H.S., J.F., K.M., E.F.P., O.M., V.C.G., supervised by T.A.C., K.K.R., G.M.R., M.B.A.

Cleaning and Analysis of Data: H.S., J.F., R.A., V.C.G., E.F.P, L.V., M.E., supervised by K.M.T., K.K.R., M.B.A.

Initial Drafting of Manuscript: H.S., K.K.R., M.B.A. Corrected and Approved Manuscript: All Authors

## Notes

### Competing Interest Statement

The authors have declared no competing interest.

