## Supplementary for "Reduced antibacterial drug resistance and *bla*_CTX-M_ β-lactamase gene carriage in cattle-associated *Escherichia coli* at low temperatures, at sites dominated by older animals and on pastureland: implications for surveillance"

**Supplementary Methods**

**Data acquisition**

**Study population**

A convenience sample representing a variety of dairy management systems was included, ranging from seasonally calving extensively managed herds to zero-grazed intensive systems. Dairy farms in this dataset were comparable to farms throughout the UK, with a median herd size of 193 compared to a UK median of 178, a median 305-day milk yield of 7488 L compared to a UK median of 8967 L, and a median somatic cell count (SCC) of 167,000 cells/mL of milk compared to a UK median of 178,000 cells/mL of milk (https://www.nmr.co.uk/uploads/files/files/HolsteinFriesian-500Herds-Aug2018(1).pdf). Antibiotic purchasing in 2016 was 26 milligrams per population corrected unit (mg/PCU; European Surveillance of Veterinary Antimicrobial Consumption) for the UK dairy industry and 21 mg/PCU for represented farms (UK Veterinary Antibiotic Resistance and Sales Surveillance, 2015).

**Samples collected from representative areas**

1. The faecally contaminated environment of the milking cow (Adult) population - either the collecting yard, housing shed or field, collected every month from 52 farms (one farm was a youngstock-only unit). A total of 1963 of samples of this type were collected.
2. The faecally contaminated environment of the dry cow population (Dry Cow), collected from 46 farms. A total of 321 of samples of this type were collected.
3. The faecally contaminated environment of replacement dairy heifers. On 41 farms, approximately 10 heifers were followed from birth until 18 months of age, with samples collected monthly from their environment. In addition, 10 further farms had samples collected from pre-weaned calves (calves still being fed milk) to increase the number of samples available for this group. For analysis, these samples were divided into pre-weaned calves (671 Calf samples) and post-weaned heifers (1337 Heifer samples) due to the differing management of these two groups.
4. Public footpaths or other rights of way that crossed the farm, where relevant (Footpath). A total of 547 samples of this type were collected from 40 farms.

**Management Questionnaires**

*Questionnaire 1* (**Figure S3**) was completed by the researcher in the presence of the farmer at the time of consent and the first farm visit

*Questionnaire 2* was completed by the researcher in the presence of the farmer using Epicollect5 approximately six to nine months into the project. The project can be viewed at

https://five.epicollect.net/project/ohstar-q2.

*Questionnaire 3* was completed by the researcher in the presence of the farmer using Epicollect5 approximately 12-16 months into the project. Details can be viewed at https://five.epicollect.net/project/ohstar-q3.

*Questionnaire 4* (**Figure S4**) was completed by the researcher during a telephone call with the farmer within two months of the last visit to the farm.

**Questionnaire validation and processing**

In total 610 variables were derived from the four questionnaires. These were validated and processed in the following way.

1. Variables with a single response or zero variance were removed.
2. Variables that had either been shown in the literature to be important risk factors or which were judged by veterinary experts to be of potential importance were selected.
3. Categorical levels were collapsed to avoid small response counts.
4. Some variables were combined as individually they provided detail irrelevant to this study.
5. Repeat questions used to derive demographic variables (from Questionnaires 1 to 4) were averaged.
6. Variables with missing values were removed; all variables removed for this reason were also considered to likely be of low importance

**Sample characteristics**

Information for each sample was recorded on a datasheet at the time of collection. Different information was recorded depending on the sample type.

*All samples:* sample number; textual description of sample location; whether animals were housed or in a field; OS reference for outdoor samples

*Pre-weaned calf environment samples:* sample number; eartag numbers; textual description of the sample location; total number of animals in the group; presence or absence of beef calves in the group; date of birth for each calf dry cow therapy used on the dam of each calf in the dry period before the birth

*Footpaths*: presence of livestock at time of sampling

**Antimicrobial usage**

All dairy farmer participants gave permission for researchers to contact their veterinary practices and request antimicrobial prescription/sales data for a period of at least one year before the beginning of the project through the end of the project.

All practices except one supplied records. This practice serviced two farms and on-farm records were used instead for these two farms.

Data were assessed by a veterinary researcher to ensure consistent naming of products and quantification between practices, given a wide range of variations on product names and quantity denominators used.

Usage metrics were produced using mg/PCU (European Surveillance of Veterinary Antimicrobial Consumption) for the first 12 months of the project, from the date the farm enrolled on the project and had the first samples collected (range January 2017- July 2017) until 12 months after this date (range January 2018-July 2018). Variables were created for:

- Total usage of all antibiotics
- Total usage of first-generation cephalosporins
- Total usage of third- and fourth-generation cephalosporins

**Data analysis**

**Modelling**

All code can be found at https://github.com/HannahSchubert1/OH-STAR-modelling-code.

***Modelling risk of bla_CTX-M_ β-lactamase gene carriage***

Risk factor analysis was performed separately on Calf data and on the full dataset (all samples combined). Thirty-seven variables were selected for the Calf risk factor analysis and 110 for the full risk factor analysis as described below. Exploratory data analysis revealed the main source of clustering of observations was at the farm level. There were no noticeable longitudinal patterns or clustering due to the location within each farm. Random intercept logistic regression model with farm as a random effect were used throughout the analysis.

Two approaches to risk factor analysis were performed: a variable selection method (using univariable screening followed by step-down model selection; method 1) and a Bayesian method with regularization; method 2 (on the full dataset only).

*Method 1*

**Variable selection**

Variable selection was performed by first screening the variables for association with *E. coli* carrying *bla*_CTX-M_ using univariable, multilevel logistic regression (Dohoo 2011; Veterinary Epidemiologic Research, 2^nd^ Edition) with each variable entered as a fixed effect and with random intercepts for each farm. For the analysis of the full dataset, it was not possible to converge such a model for 27 variables and so 83 variables remained, most likely due to the low number of samples that were positive for *E. coli* carrying *bla*_CTX-M_. See **Tables S1** and **S2** for full univariable results.

Variables with associations where p <0.25 after controlling for false discovery rate using the Benjamini-Hochberg procedure (chosen to be the most appropriate method for the exploratory nature of the project) were entered in to a multivariable, multilevel logistic regression model with random intercepts for each farm. A backwards stepwise procedure was used to further refine the model, selecting only those variables where the regression coefficient maintained p<0.05. Variables which survived this analysis were checked for multicollinearity by removing each variable each in turn and checking that the confidence intervals for the estimate for each variable still overlapped. The predictive accuracy was checked using area under the Receiver Operating Characteristic Curve (0.84 for the full dataset; 0.80 for Calf data).

**Accounting for measurement error**

In order to account for measurement error of the *bla*_CTX-M_-positive *E. coli* (whereby if more *E. coli* are found in a sample, the sensitivity of the test for finding *bla*_CTX-M_-positive *E. coli* is higher) the logistic link function was altered to include the sensitivity and specificity of each sample, following the work by Coutinho et al. (Coutinho C, Bastos LS, Corrêa da Mota J, Toledo L, Costa K, Bertoni N, Bastos FI; The risks of HCV infection among Brazilian crack cocaine users: incorporating diagnostic test uncertainty Sci Rep. 2019; 9: 443). The altered logistic link function was derived by equating the conditional probability of the true *bla*_CTX-M_ status $P(Y^{true}=1|X)$ to the conditional probability of the observed *bla*_CTX-M_ status, $P(Y^{obs}=1|X)$ for a given sensitivity (s) and specificity (e). The R-code for this link function is given in full below.

The specificity of the *bla*_CTX-M_ test was assumed to be 100% across samples whereas the sensitivity ($s)$, was estimated separately for each observation using the number of *E. coli* colonies grown on antibiotic-free agar $(k)$ and a minimum detectable prevalence of *bla*_CTX-M_-positive *E. coli* $(q)$ through:

$$s=1-(1-q)^{k}$$

The distribution $s$ across the whole dataset for different values of $q$ is shown in **Figure S5**. While the value of $q$ did change the sensitivity associated with each observation, it had only a negligible effect on the coefficient estimates (**Fig S2A**) and AUC (**Fig S2B**) over the range $0.01 - 1.00$. The reported values used a $q$ of 0.01.

logitse <- function(s=1, e=1)
 {

 linkfun <- function(mu) qlogis((mu+e-1)/(s+e-1))

 linkinv <- function(eta) plogis(eta) * (s+e-1) + 1 - e

 logit_mu_eta <- function(eta) {
 ifelse(abs(eta)>30,.Machine$double.eps,
 exp(eta)/(1+exp(eta))^2)
 }

 mu.eta <- function(eta) (s + e - 1) *logit_mu_eta(eta)

 valideta <- function(eta) TRUE

 link <- paste("logitse(",deparse(substitute(s)),",",
 deparse(substitute(e)), ")", sep="")

 structure(list(linkfun = linkfun,
 linkinv = linkinv,
 mu.eta = mu.eta,
 logit_mu_eta = logit_mu_eta,
 valideta = valideta,
 name = link),
 class = "link-glm")
}

*Method 2:* **Bayesian model for predicting presence of *bla*_CTX-M_-positive *E. coli***

### A farm-level random intercept model was fit using the R package BRMS on the full dataset. All variables were used as predictors of *bla*_CTX-M_-positive *E. coli* but were split into two groups. The first group comprised farm-level 1) total antibiotic use, 2) total third- and fourth-generation cephalosporin use, and 3) total first-generation cephalosporin use. The second group contained the remaining meteorological and farm management variables. For the first group, uninformative priors (normal distribution with a mean 0, standard deviation 5) were used; for the second group, a regularizing prior (horseshoe prior with a single degree of freedom [Carvalho CM, Polson NG, Scott JG, The horseshoe estimator for sparse signals, Biometrika, 2010; 97:465-480]) was used. The mean intercept was also given a diffuse prior (normal distribution with mean 0, standard deviation 5) while the standard deviation of the random effects was given a half-Student-T distribution with three degrees of freedom and a scale factor of 10 (the default in BRMS). Four independent Markov chains were sampled with 1000 warmup iterations and 1000 sampling iterations. The target acceptance criterion was increased from the default 0.8 to 0.95 to decrease the chance of sampling divergencies (although two remained, these were not deemed significant). The sampling was assumed to be well converged as the Gelman-Rubin statistic for each variable was 1.00. Results are shown in Table S3.

***Model for predicting presence of E. coli resistant to non-cephalosporins***

For each resistance phenotype – amoxicillin, ciprofloxacin, streptomycin and tetracycline, a random intercept model with farm as the random effect was fitted using the R package BRMS on the full dataset (https://cran.r-project.org/web/packages/brms/index.html). All variables (see above) were used as predictors of resistant *E. coli* in a sample but were split into two groups. The first group (the ‘main’ variables) comprised farm-level ABU and average monthly temperature at the time of sample collection as these were the main variables hypothesised to influence resistance. For all models total ABU, streptomycin usage, tetracycline usage, amoxicillin usage, fluoroquinolone usage and cefalexin usage were included as predictors. For each model, the usage of additional antibacterial drugs was tested if they were hypothesised to be important in selecting for resistance to the relevant model: for amoxicillin model, first-generation cephalosporins, penicillins, potentiated amoxicillins; ciprofloxacin model, novobiocin, third- and fourth-generation cephalosporins; streptomycin and tetracycline models, no additional ABU variables were tested.

The second group of variables (the ‘regularised’ variables) contained the remaining farm management variables.

For the first group (the ‘main’ variables), uninformative priors (normal distribution with a mean 0, standard deviation 5) were used; for the second group (the ‘regularised’ variables), a regularising prior (horseshoe prior with a single degree of freedom) was used. The mean intercept was also given a diffuse prior (normal distribution with mean 0, standard deviation 5) while the standard deviation of the random effects was given a half-Student-T distribution with three degrees of freedom and a scale factor of 10 (the default in BRMS). Four independent Markov chains were sampled with 1,000 warmup iterations and 10,000 sampling iterations. The target acceptance criterion was increased from the default 0.8 to 0.98 to decrease the chance of sampling divergencies in all models. Measurement error based on *E. coli* density was accounted for as described above. All reported results used a q of 0.01.

In addition, the measurement error of temperature was considered because only an average monthly temperature was available. Data where daily temperature was recorded for a (different) twelve-month period were used to calculate an estimated average monthly standard deviation over the twelve months (2.89). The ‘me’ function in the package ‘BRMS’ (https://rdrr.io/cran/brms/man/me.html) was used, whereby the monthly temperature variable was assumed to vary with a standard deviation of 2.89 (0.61 when scaled as the temperature was scaled to mean 0 and standard deviation 1 before entering the model; the actual standard deviation of the temperature was 4.7).

To further test the robustness of the model outputs, all models were re-run using sceptical priors, whereby the prior was set to the opposite of what would be expected. So, for all main variables, if there was an association, a positive association would be expected given prior knowledge. To test the models with sceptical priors, a prior for the main variables was given as a normal distribution with mean -0.5 and a narrow standard deviation 2 (i.e. assuming a negative correlation with low variation; the opposite of expected).

**Bayesian model checking**

Convergence is assumed to be good with a Gelman-rubin statistic (Rhat) of 1.00 for all variables across all models. **Figure S6** shows trace plots for the associated variables, also providing evidence of good convergence.

There were no divergence issues reported for any of the models.

**Table S4** shows full results from all models, including odds ratios, 95% credible intervals and effective sample size.

**Figure S5** shows the posterior distributions of the associated variables.

Re-running the models with sceptical priors did not alter the model conclusions (table S2).

**Supplementary Tables**

**Table S1:** All univariable analyses from the Calf dataset. All variables presented in the table were screened for association with *E. coli* carrying *bla*_CTX-M_ carriage using univariable, multilevel logistic regression with each variable entered as a fixed effect and with random intercepts for each farm. Variables with associations where the adjusted p <0.25 (after controlling for false discovery rate using the Benjamini-Hochberg procedure) were taken forward into the backwards stepwise model.

| **Variable** | **Odds ratio** | **Lower CI** | **Upper CI** | **p** | **Adjusted p** | **Description** |
| --- | --- | --- | --- | --- | --- | --- |
| s_temp | 1.79 | 1.39 | 2.30 | 0.00 | 0.00 | Average monthly temperature |
| s_cefq_dct_6m | 4.22 | 1.80 | 9.90 | 0.00 | 0.01 | Use of cefquinome dry cow therapy in the last six months |
| s_fram_dct_6m | 2.95 | 1.45 | 5.98 | 0.00 | 0.03 | Use of framycetin dry cow therapy in the last six months |
| f_trough_clean | 0.43 | 0.24 | 0.77 | 0.00 | 0.04 | Daily cleaning of water troughs in calf housing |
| f_nsaiddiarr | 0.28 | 0.10 | 0.76 | 0.01 | 0.08 | Routine use of anti-inflammatories in cases of calf diarrhoea |
| f_ceph_1 | 0.45 | 0.23 | 0.88 | 0.02 | 0.12 | Total amount of first-generation cephalosporin use |
| f_poultry | 2.64 | 1.05 | 6.63 | 0.04 | 0.19 | Presence of poultry on the farm |
| f_rain | 1.18 | 0.93 | 1.49 | 0.17 | 0.62 | Average monthly rainfall |
| f_give_col | 2.01 | 0.72 | 5.58 | 0.18 | 0.62 | Administration of colostrum within six hours of life |
| f_pneum_vacc | 0.51 | 0.20 | 1.29 | 0.15 | 0.62 | Routine use of calf respiratory vaccination |
| f_calf_housing_type | 0.52 | 0.16 | 1.67 | 0.27 | 0.76 | Type of calf housing |
| f_anticocc | 1.68 | 0.68 | 4.13 | 0.26 | 0.76 | Routine use of anticoccidials |
| f_calving_group | 1.85 | 0.41 | 8.40 | 0.43 | 0.80 | Whether calves were born in a group pen |
| s_clox_dct_6m | 0.62 | 0.22 | 1.76 | 0.37 | 0.80 | Use of cloxacillin dry cow therapy in the last six months |
| f_scc | 0.84 | 0.55 | 1.28 | 0.41 | 0.80 | Average herd somatic cell count |
| f_geographical_area | 1.66 | 0.49 | 5.61 | 0.42 | 0.80 | Geographical location of the farm |
| f_heifers_waste | 1.44 | 0.59 | 3.53 | 0.42 | 0.80 | Whether waste milk has been fed to heifers |
| f_anything_waste | 1.45 | 0.54 | 3.88 | 0.46 | 0.83 | Whether waste milk was fed to any calves |
| f_pattern | 1.50 | 0.44 | 5.09 | 0.52 | 0.86 | Calving pattern |
| f_water | 1.32 | 0.49 | 3.59 | 0.58 | 0.86 | Source of drinking water |
| f_calf_housing_older | 0.82 | 0.28 | 2.40 | 0.71 | 0.86 | Whether calves have been kept near to older animals |
| s_ceph_dct_6m | 1.18 | 0.48 | 2.95 | 0.72 | 0.86 | Use of cephalonium dry cow therapy in the last six months |
| f_um_spray | 0.72 | 0.22 | 2.36 | 0.59 | 0.86 | Routine use of umbilical treatment |
| f_time_dam | 0.86 | 0.45 | 1.62 | 0.63 | 0.86 | Amount of time spent with dam |
| f_halocur | 1.16 | 0.46 | 2.98 | 0.75 | 0.86 | Routine use of treatment to prevent cryptosporidiosis |
| f_diarrvacc | 0.79 | 0.30 | 2.06 | 0.63 | 0.86 | Routine use of calf enteric disease vaccination |
| f_bought_pre | 0.92 | 0.61 | 1.39 | 0.70 | 0.86 | Number of cattle bought in the 12 months before the start of the project |
| f_herd_size | 1.08 | 0.70 | 1.67 | 0.73 | 0.86 | Number of milking cows on the farm |
| f_yield | 1.08 | 0.69 | 1.69 | 0.72 | 0.86 | Average herd yield |
| f_total_cattle | 0.92 | 0.59 | 1.42 | 0.69 | 0.86 | Total number of cattle on the farm |
| f_total_mg_pcu | 1.04 | 0.70 | 1.54 | 0.84 | 0.90 | Total antibiotic usage |
| f_wean | 1.16 | 0.34 | 3.93 | 0.81 | 0.90 | Age of weaning |
| f_equine | 0.94 | 0.39 | 2.30 | 0.90 | 0.92 | Presence of horses on the farm |
| f_ceph_3_4 | 1.02 | 0.64 | 1.64 | 0.93 | 0.93 | Amount of third- and fourth-generation cephalosporins used |

**Table S2**: All univariable analyses from the full dataset. All variables presented in the table were screened for association with *E. coli* carrying *bla*_CTX-M_ carriage using univariable, multilevel logistic regression with each variable entered as a fixed effect and with random intercepts for each farm. Variables with associations where the adjusted p <0.25 (after controlling for false discovery rate using the Benjamini-Hochberg procedure) were taken forwards into the backwards stepwise model.

| **Variable** | **Odds ratio** | **Lower CI** | **Upper CI** | **p** | **Adjusted p** | **Description** |
| --- | --- | --- | --- | --- | --- | --- |
| preweaned_heifer | 5.85 | 4.33 | 7.89 | 0.00 | 0.00 | Sample collected from a pre-weaned heifer |
| s_temp | 1.52 | 1.32 | 1.76 | 0.00 | 0.00 | Average monthly temperature |
| weaned_heifer | 0.48 | 0.33 | 0.68 | 0.00 | 0.00 | Sample collected from a weaned heifer |
| s_housed_outdoor | 0.30 | 0.17 | 0.54 | 0.00 | 0.00 | Sample collected from pastureland |
| f_foragetype_3 | 3.11 | 1.51 | 6.43 | 0.00 | 0.03 | Maize silage used on farm |
| footpath | 0.37 | 0.20 | 0.70 | 0.00 | 0.03 | Sample collected from a public footpath |
| adult | 0.62 | 0.46 | 0.84 | 0.00 | 0.03 | Sample collected from a milking cow |
| dry | 0.40 | 0.20 | 0.81 | 0.01 | 0.12 | Sample collected from a dry cow |
| f_othergraze | 0.47 | 0.25 | 0.86 | 0.02 | 0.15 | Use of grazing land away from the farm |
| f_salmvacc | 0.15 | 0.03 | 0.83 | 0.03 | 0.24 | Routine vaccination against Salmonella |
| f_trough_clean | 0.67 | 0.43 | 1.04 | 0.07 | 0.52 | Daily cleaning of water troughs in calf housing |
| f_cefq_dct | 1.90 | 0.90 | 4.01 | 0.09 | 0.59 | Use of cefquinome dry cow therapy |
| f_acf | 1.71 | 0.90 | 3.21 | 0.10 | 0.59 | Use of automatic cluster flushing |
| f_scc | 0.75 | 0.53 | 1.06 | 0.10 | 0.59 | Average herd somatic cell count |
| f_total_mg_pcu | 1.23 | 0.94 | 1.60 | 0.13 | 0.62 | Total antibiotic usage |
| f_manure_7 | 0.14 | 0.01 | 1.67 | 0.12 | 0.62 | Use of composter for muck management |
| f_manure_1 | 1.61 | 0.84 | 3.07 | 0.15 | 0.62 | Use of slurry lagoon |
| f_hunt | 0.61 | 0.32 | 1.17 | 0.14 | 0.62 | Whether the hunt crosses the farm |
| f_um_spray | 0.54 | 0.23 | 1.25 | 0.15 | 0.62 | Routine use of umbilical treatment |
| f_anticocc | 1.60 | 0.84 | 3.05 | 0.15 | 0.62 | Routine use of anticoccidials |
| f_lungvacc | 0.63 | 0.31 | 1.25 | 0.18 | 0.64 | Routine vaccination against lungworm |
| f_bvdvacc | 1.59 | 0.79 | 3.19 | 0.19 | 0.64 | Routine vaccination against bovine viral diarrhoea |
| f_give_col | 1.58 | 0.79 | 3.16 | 0.19 | 0.64 | Administration of colostrum within six hours of life |
| f_time_dam | 0.74 | 0.47 | 1.16 | 0.19 | 0.64 | Time calf spends with the dam |
| f_poultry | 1.56 | 0.82 | 2.97 | 0.18 | 0.64 | Presence of poultry on the farm |
| f_manure_3 | 0.66 | 0.32 | 1.37 | 0.27 | 0.73 | Use of a metal slurry store |
| f_bedmilk_6 | 3.39 | 0.46 | 25.07 | 0.23 | 0.73 | Use of green bedding for milking cows |
| f_rat | 1.47 | 0.75 | 2.89 | 0.27 | 0.73 | Perceived level of rat infestation on the farm |
| f_cleancluster | 1.27 | 0.82 | 1.98 | 0.28 | 0.73 | How often the milking cluster was cleaned |
| f_timesfirstmast | 1.18 | 0.87 | 1.61 | 0.28 | 0.73 | How often clinical mastitis cases were treated with intramammary tubes |
| f_leptovacc | 1.50 | 0.71 | 3.16 | 0.29 | 0.73 | Routine use of leptospirosis vaccination |
| f_nsaiddiarr | 0.64 | 0.30 | 1.36 | 0.24 | 0.73 | Routine use of anti-inflammatories in the case of calf diarrhoea |
| f_anything_waste | 1.49 | 0.74 | 2.98 | 0.26 | 0.73 | Whether waste milk was fed to any calves |
| f_halocur | 1.42 | 0.73 | 2.77 | 0.31 | 0.75 | Routine use of treatment to prevent cryptosporidiosis |
| f_water | 1.39 | 0.69 | 2.81 | 0.36 | 0.84 | Source of drinking water |
| f_bedmilk_2 | 1.36 | 0.69 | 2.65 | 0.37 | 0.85 | Use of straw as milking cow bedding |
| f_manure_8 | 1.47 | 0.56 | 3.85 | 0.43 | 0.88 | Use of separator for muck management |
| f_manure_2 | 1.36 | 0.58 | 3.19 | 0.47 | 0.88 | Use of a weeping wall for slurry management |
| f_bedmilk_3 | 0.78 | 0.41 | 1.50 | 0.46 | 0.88 | Use of sawdust as milking cow bedding |
| f_starling | 0.77 | 0.39 | 1.52 | 0.46 | 0.88 | Perceived amount of starling infestation |
| f_abcalve | 1.42 | 0.62 | 3.23 | 0.40 | 0.88 | Routine use of antibiotics following a difficult calving |
| f_organic | 0.58 | 0.14 | 2.48 | 0.47 | 0.88 | Whether the farm is certified organic |
| f_clostvacc | 1.34 | 0.60 | 2.99 | 0.48 | 0.88 | Routine use of clostridial vaccine |
| s_rain | 0.95 | 0.83 | 1.09 | 0.48 | 0.88 | Average monthly rainfall |
| f_manure_4 | 0.81 | 0.43 | 1.54 | 0.52 | 0.89 | Use of a concrete slurry store |
| f_foragetype_5 | 1.21 | 0.59 | 2.45 | 0.61 | 0.89 | Feeding of hay |
| f_foragetype_2 | 0.77 | 0.29 | 2.06 | 0.61 | 0.89 | Feeding of big bale grass silage |
| f_aiexternal | 1.21 | 0.64 | 2.30 | 0.55 | 0.89 | Artificial insemination performed by external company |
| f_pheasant | 0.80 | 0.33 | 1.91 | 0.61 | 0.89 | Perceived amount of pheasants on farmland |
| f_deer | 1.26 | 0.59 | 2.69 | 0.55 | 0.89 | Perceived amount of deer on farmland |
| f_shows | 1.47 | 0.46 | 4.72 | 0.52 | 0.89 | Whether cattle go to shows |
| f_nsaidmast | 1.09 | 0.81 | 1.47 | 0.58 | 0.89 | Routine use of anti-inflammatories in cases of mastitis |
| f_calf_housing_older | 0.81 | 0.37 | 1.78 | 0.60 | 0.89 | Whether calves have been kept near to older animals |
| f_herd_size | 1.10 | 0.80 | 1.51 | 0.56 | 0.89 | Number of milking cows on the farm |
| f_heifers_waste | 1.20 | 0.62 | 2.35 | 0.59 | 0.89 | Whether waste milk has been fed to heifers |
| f_machinery | 0.86 | 0.45 | 1.63 | 0.64 | 0.90 | Whether machinery was shared with other farms |
| f_muckspreader | 0.86 | 0.45 | 1.66 | 0.66 | 0.90 | Use of a contract muck spreader |
| f_calving_group | 1.30 | 0.44 | 3.82 | 0.64 | 0.90 | Whether calves were born in a group pen |
| f_diarrvacc | 0.85 | 0.43 | 1.69 | 0.65 | 0.90 | Routine use of calf enteric disease vaccination |
| s_calving_now | 1.15 | 0.60 | 2.22 | 0.67 | 0.90 | Whether cows were calving during the month in which the sample was taken |
| f_ceph_1 | 0.97 | 0.72 | 1.32 | 0.85 | 0.93 | Total amount of first-generation cephalosporin use |
| f_ceph_3_4 | 0.95 | 0.70 | 1.29 | 0.75 | 0.93 | Amount of third- and fourth- generation cephalosporins used |
| f_clox_dct | 0.92 | 0.42 | 2.02 | 0.83 | 0.93 | Use of cloxacillin dry cow therapy |
| f_foragetype_4 | 1.12 | 0.59 | 2.13 | 0.73 | 0.93 | Feeding of whole crop silage |
| f_bedmilk_4 | 0.66 | 0.06 | 7.33 | 0.74 | 0.93 | Use of paper waste for milking cow bedding |
| f_shoot | 1.09 | 0.57 | 2.06 | 0.80 | 0.93 | Whether the shoot crosses the farm |
| f_badger | 1.10 | 0.40 | 3.01 | 0.86 | 0.93 | Perceived badger population on the farm |
| f_premises | 0.94 | 0.49 | 1.81 | 0.86 | 0.93 | Number of farm premises |
| f_drytrough | 0.95 | 0.60 | 1.50 | 0.84 | 0.93 | How often the dry cow water trough was cleaned out |
| f_treatfoot | 1.07 | 0.57 | 2.01 | 0.84 | 0.93 | Use of a footbath for lameness outbreaks |
| f_daystreatmast | 1.04 | 0.76 | 1.43 | 0.81 | 0.93 | Number of days cases of mastitis were treated with antibiotics |
| f_hectares | 1.03 | 0.75 | 1.42 | 0.84 | 0.93 | Size of farm in hectares |
| f_wean | 0.88 | 0.33 | 2.31 | 0.79 | 0.93 | Age of weaning |
| f_equine | 0.94 | 0.49 | 1.79 | 0.85 | 0.93 | Presence of horses on the farm |
| f_manure_6 | 0.97 | 0.51 | 1.84 | 0.93 | 0.96 | Use of muck heap on concrete |
| f_feedlorry | 0.97 | 0.49 | 1.91 | 0.92 | 0.96 | Whether feed lorries crossed any cattle yards when delivering |
| f_lime | 1.03 | 0.53 | 1.99 | 0.93 | 0.96 | Use of lime in milking cow housing |
| f_pneum_vacc | 0.97 | 0.49 | 1.92 | 0.93 | 0.96 | Routine use of calf respiratory vaccination |
| f_total_cattle | 0.98 | 0.71 | 1.36 | 0.92 | 0.96 | Total number of cattle on the farm |
| f_foragetype_1 | 1.04 | 0.25 | 4.26 | 0.96 | 0.97 | Feeding of clamp grass silage |
| f_bedmilk_1 | 0.99 | 0.49 | 2.00 | 0.99 | 0.99 | Use of sand in milking cow bedding |

**Table S3:** Results of the Bayesian analysis, odds ratios with 95% credible intervals. The type of prior (diffuse/regularized) is also shown. Variables in *italics* were regularized and are ordered according to the average (for factor variables with more than two levels) magnitude of their effect.

| **Variable** | **Odds ratio** | **Lower CI** | **Upper CI** | **Description** |
| --- | --- | --- | --- | --- |
| total_mg_pcu | 1.16 | 0.78 | 1.69 | Total antibiotic usage |
| ceph_1 | 1.01 | 0.69 | 1.45 | Total amount of first-generation cephalosporin use |
| ceph_3_4 | 0.96 | 0.65 | 1.45 | Amount of third- and fourth- generation cephalosporins used |
| *dry1* | 0.96 | 0.63 | 1.17 | Sample collected from the dry cow environment |
| *f_abcalve1* | 1.02 | 0.74 | 1.59 | Routine use of antibiotics following a difficult calving |
| *f_acf1* | 1.09 | 0.85 | 2.13 | Use of automatic cluster flushing |
| *f_aiexternal1* | 1.07 | 0.87 | 1.85 | Artificial insemination performed by external company |
| *f_anticocc1* | 1.04 | 0.81 | 1.61 | Routine use of anticoccidials |
| *f_anything_waste1* | 1.03 | 0.81 | 1.59 | Whether waste milk was fed to any calves |
| *f_badger2* | 1.01 | 0.74 | 1.51 | Perceived badger population on the farm |
| *f_bedmilk_1yes* | 0.99 | 0.67 | 1.37 | Use of sand in milking cow bedding |
| *f_bedmilk_2yes* | 1.03 | 0.79 | 1.54 | Use of straw as milking cow bedding |
| *f_bedmilk_3yes* | 0.98 | 0.67 | 1.24 | Use of sawdust as milking cow bedding |
| *f_bedmilk_4yes* | 1.00 | 0.60 | 1.60 | Use of paper waste for milking cow bedding |
| *f_bedmilk_6yes* | 1.02 | 0.68 | 1.83 | Use of green bedding for milking cows |
| *f_bought_pre* | 0.96 | 0.69 | 1.12 | Number of cattle bought in the 12 months before the start of the project |
| *f_bvdvaccY* | 1.07 | 0.84 | 1.93 | Routine vaccination against bovine viral diarrhoea |
| *f_calf_housing_older1* | 0.98 | 0.67 | 1.30 | Whether calves have been kept near to older animals |
| *f_calf_housing_type3* | 1.01 | 0.76 | 1.40 | Type of calf housing |
| *f_calveclean.L* | 1.00 | 0.76 | 1.31 | How often the calving pens are cleaned out |
| *f_calving_group1* | 1.02 | 0.74 | 1.63 | Whether calves were born in a group pen |
| *f_cefq_dct1* | 1.04 | 0.80 | 1.66 | Use of cefquinome dry cow therapy |
| *f_ceph_dct1* | 0.84 | 0.31 | 1.15 | Use of cephalonium dry cow therapy |
| *f_cleancluster.L* | 1.03 | 0.85 | 1.42 | How often the milking cluster was cleaned |
| *f_clostvaccY* | 1.01 | 0.73 | 1.48 | Routine use of clostridial vaccine |
| *f_clox_dct1* | 1.03 | 0.77 | 1.63 | Use of cloxacillin dry cow therapy |
| *f_daystreatmast* | 1.00 | 0.82 | 1.24 | Number of days cases of mastitis were treated with antibiotics |
| *f_deer2* | 1.01 | 0.74 | 1.38 | Perceived amount of deer on farmland |
| *f_diarrvaccY* | 0.97 | 0.68 | 1.22 | Routine use of calf enteric disease vaccination |
| *f_drytrough.L* | 0.98 | 0.70 | 1.20 | How often the dry cow water trough was cleaned out |
| *f_equine1* | 0.97 | 0.67 | 1.24 | Presence of horses on the farm |
| *f_feedlorry1* | 0.97 | 0.66 | 1.26 | Whether feed lorries crossed any cattle yards when delivering |
| *f_firstmastitisCobactan* | 1.02 | 0.67 | 1.71 | First line mastitis treatment |
| *f_firstmastitisMastiplanLC* | 0.92 | 0.29 | 1.34 |  |
| *f_firstmastitisOrbeninLA* | 0.93 | 0.33 | 1.41 |  |
| *f_firstmastitistdDmultiject* | 1.01 | 0.76 | 1.40 |  |
| *f_firstmastitisUbrolexin* | 1.07 | 0.80 | 2.33 |  |
| *f_firstmastitisUbroYellow* | 0.98 | 0.66 | 1.27 |  |
| *f_foragetype_1yes* | 0.98 | 0.63 | 1.38 | Feeding of clamp grass silage |
| *f_foragetype_2yes* | 0.95 | 0.51 | 1.24 | Feeding of big bale grass silage |
| *f_foragetype_3yes* | 1.32 | 0.87 | 4.36 | Maize silage used on farm |
| *f_foragetype_4yes* | 1.02 | 0.81 | 1.46 | Feeding of whole crop silage |
| *f_foragetype_5yes* | 1.02 | 0.79 | 1.56 | Feeding of hay |
| *f_foragetype_6yes* | 1.18 | 0.81 | 4.84 | Feeding of straw |
| *f_foragetype_7yes* | 0.98 | 0.69 | 1.29 | Feeding of haylage |
| *f_fox2* | 1.00 | 0.75 | 1.33 | Perceived amount of foxes on farmland |
| *f_fram_dct1* | 1.56 | 0.93 | 5.06 | Use of framycetin dry cow therapy |
| *f_geographical_area2* | 0.99 | 0.70 | 1.34 | Farm is in geographical area 2 |
| *f_geographical_area3* | 1.15 | 0.87 | 2.75 | Farm is in geographical area 3 |
| *f_give_col1* | 1.03 | 0.78 | 1.57 | Administration of colostrum within six hours of life |
| *f_halocur1* | 1.01 | 0.79 | 1.41 | Routine use of treatment to prevent cryptosporidiosis |
| *f_hectares* | 1.01 | 0.85 | 1.26 | Size of farm in hectares |
| *f_heifers_waste1* | 1.09 | 0.85 | 2.15 | Whether waste milk has been fed to heifers |
| *f_herd_size* | 1.01 | 0.83 | 1.26 | Number of milking cows on the farm |
| *f_hostfarmwalk1* | 0.97 | 0.62 | 1.23 | Whether a farm walk has taken place on the farm in the last 2 years |
| *f_hunt2* | 0.95 | 0.57 | 1.18 | Whether the hunt crosses the farm |
| *f_ibrvaccY* | 1.02 | 0.78 | 1.54 | Routine use of IBR vaccination |
| *f_injectmast* | 1.03 | 0.86 | 1.40 | Whether mastitis cases are routinely injected with antibiotic |
| *f_leptovaccY* | 1.00 | 0.74 | 1.34 | Routine use of leptospirosis vaccination |
| *f_lime1* | 0.97 | 0.65 | 1.26 | Use of lime in milking cow housing |
| *f_lungvaccY* | 0.94 | 0.53 | 1.19 | Routine vaccination against lungworm |
| *f_machinery1* | 0.96 | 0.61 | 1.22 | Whether machinery was shared with other farms |
| *f_manure_1yes* | 1.08 | 0.85 | 1.99 | Use of slurry lagoon |
| *f_manure_2yes* | 1.03 | 0.80 | 1.65 | Use of a weeping wall for slurry management |
| *f_manure_3yes* | 0.97 | 0.65 | 1.25 | Use of a metal slurry store |
| *f_manure_4yes* | 0.98 | 0.69 | 1.26 | Use of a concrete slurry store |
| *f_manure_5yes* | 0.97 | 0.61 | 1.23 | Muck heap in field used |
| *f_manure_6yes* | 0.99 | 0.71 | 1.28 | Use of muck heap on concrete |
| *f_manure_7yes* | 0.92 | 0.32 | 1.35 | Use of composter for muck management |
| *f_manure_8yes* | 1.03 | 0.74 | 1.67 | Use of separator for muck management |
| *f_market1* | 1.01 | 0.76 | 1.44 | Whether cattle are taken to market regularly |
| *f_milkgraze1* | 1.02 | 0.76 | 1.58 | Whether milking cows graze |
| *f_muckspreader3* | 1.04 | 0.72 | 2.02 | Use of a contract muck spreader |
| *f_nsaidcalve1* | 0.98 | 0.69 | 1.26 | Routine use of anti-inflammatories at calving |
| *f_nsaiddiarr1* | 0.97 | 0.62 | 1.30 | Routine use of anti-inflammatories in cases of calf diarrhoea |
| *f_nsaidmast* | 0.99 | 0.80 | 1.18 | Routine use of anti-inflammatories in cases of mastitis |
| *f_organic1* | 1.01 | 0.68 | 1.62 | Whether the farm is certified organic |
| *f_othergrazeY* | 0.91 | 0.44 | 1.16 | Use of grazing land away from the farm |
| *f_outsource1* | 0.86 | 0.26 | 1.23 | Whether heifer calves are reared by a different enterprise |
| *f_pattern2* | 1.12 | 0.84 | 2.66 | Calving pattern |
| *f_pheasant2* | 0.99 | 0.69 | 1.38 | Perceived amount of pheasants on farmland |
| *f_pigeon2* | 0.96 | 0.59 | 1.20 | Perceived level of pigeon infestation |
| *f_pneum_vacc1* | 1.00 | 0.73 | 1.37 | Routine use of calf respiratory vaccination |
| *f_poultry1* | 1.03 | 0.82 | 1.55 | Presence of poultry on the farm |
| *f_predip1* | 1.05 | 0.84 | 1.66 | Whether predip is used when milking |
| *f_premises2* | 1.01 | 0.74 | 1.45 | Number of farm premises |
| *f_rat2* | 1.02 | 0.79 | 1.43 | Perceived level of rat infestation on the farm |
| *f_rook2* | 0.94 | 0.52 | 1.20 | Perceived level of rook infestation on farm |
| *f_salmvaccY* | 0.85 | 0.20 | 1.23 | Routine vaccination against Salmonella |
| *f_scc* | 0.96 | 0.71 | 1.14 | Average herd somatic cell count |
| *f_shoot2* | 0.99 | 0.70 | 1.29 | Whether the shoot crosses the farm |
| *f_shows1* | 1.14 | 0.82 | 3.75 | Whether cattle from the farm are taken to shows |
| *f_starling2* | 0.98 | 0.69 | 1.26 | Perceived amount of starling infestation |
| *f_time_dam.L* | 0.99 | 0.76 | 1.25 | Time calf spends with dam |
| *f_timesfirstmast* | 1.02 | 0.88 | 1.28 | How often clinical mastitis cases were treated with intramammary tubes |
| *f_total_cattle* | 0.99 | 0.78 | 1.20 | Total number of cattle on the farm |
| *f_treatfoot1* | 0.99 | 0.72 | 1.31 | Use of a footbath for lameness outbreaks |
| *f_trough_clean.L* | 0.88 | 0.47 | 1.10 | Daily cleaning of water troughs in calf housing |
| *f_um_spray1* | 0.98 | 0.64 | 1.33 | Routine use of umbilical treatment |
| *f_water1* | 1.07 | 0.84 | 2.07 | Source of drinking water |
| *f_wean2* | 0.95 | 0.51 | 1.25 | Age of weaning |
| *f_wean3* | 1.12 | 0.86 | 2.48 |  |
| *f_whichclinrespMacrolide* | 0.99 | 0.69 | 1.30 | Most commonly used antibiotic to treat calf pneumonia |
| *f_whichclinrespOtherDdontknow* | 0.92 | 0.32 | 1.47 |  |
| *f_whichclinrespPenicillimoxycillin* | 0.93 | 0.31 | 1.39 |  |
| *f_whichclinrespTetracycline* | 1.00 | 0.71 | 1.40 |  |
| *f_yield* | 1.02 | 0.85 | 1.32 | Average herd yield |
| *footpath1* | 0.90 | 0.41 | 1.16 | Sample collected from a public footpath |
| *preweaned_heifer1* | 5.58 | 3.86 | 8.49 | Sample collected from the environment of a pre-weaned heifer |
| *s_calving_now1* | 1.00 | 0.74 | 1.32 | Whether cows were calving during the month in which the sample was taken |
| *s_housed_outdooroutdoor* | 0.50 | 0.22 | 1.02 | Sample collected from pastureland |
| *s_rain* | 1.19 | 0.99 | 1.47 | Average monthly rainfall |
| *s_temp* | 1.71 | 1.42 | 2.05 | Average monthly temperature |
| *weaned_heifer1* | 0.99 | 0.78 | 1.24 | Sample collected from the environment of a weaned heifer |
| *adult1* | 1.06 | 0.90 | 1.54 | Sample collected from the milking cow environment |

**Table S4:** Full details from Bayesian logistic regression models for non-cephalosporin resistance with odds ratios, 95% credible intervals and effective sample sizes. Tables are ordered by magnitude of effect size of the variables. A prefix ‘main_’ indicates variables tested as the main variables with diffuse priors. A prefix ‘reg_’ indicated regularised variables where a horseshoe prior was applied.

**Amoxicillin**

| **Variable** | **Odds ratio** | **Lower CI** | **Upper CI** | **Description** | **Effective sample size** |
| --- | --- | --- | --- | --- | --- |
| reg_s_housed_outdooroutdoor | 0.27 | 0.2 | 0.37 | Sample collected from pasture | 16776 |
| reg_preweaned_heifer1 | 1.99 | 1.29 | 2.98 | Sample collected from the environment of a pre-weaned heifer | 14162 |
| main_s_temp | 1.91 | 1.57 | 2.35 | Average monthly temperature | 14045 |
| main_total_mg_pcu | 1.61 | 0.91 | 2.86 | Total antibiotic usage | 11829 |
| main_pen | 0.71 | 0.31 | 1.67 | Total usage of penicillin | 11046 |
| main_co_amox | 1.24 | 0.82 | 1.92 | Total usage of amoxicillin-clavulonic acid | 12507 |
| reg_f_shows1 | 1.17 | 0.92 | 3.31 | Whether animals are taken to shows | 5518 |
| reg_f_firstmastitistdDmultiject | 1.17 | 0.95 | 2.22 | First line choice of mastitis treatment is streptomycin/ neomycin/ novobiocin/ penicillin | 4957 |
| main_strep | 0.88 | 0.7 | 1.11 | Total streptomycin usage | 12932 |
| reg_f_lungvaccY | 0.89 | 0.54 | 1.06 | Routine use of lungworm vaccination | 8244 |
| reg_f_foragetype_2yes | 0.9 | 0.43 | 1.08 | Feeding of big bale grass silage | 8200 |
| main_ceph_1 | 0.91 | 0.63 | 1.31 | Total usage of first generation cephalosporins | 13338 |
| main_fq | 1.1 | 0.85 | 1.43 | Total fluoroquinolone usage | 12700 |
| main_amox | 1.08 | 0.78 | 1.47 | Total amoxicillin usage | 12016 |
| reg_f_manure_4yes | 0.93 | 0.61 | 1.06 | Use of a concrete slurry store | 10621 |
| main_tet | 1.07 | 0.82 | 1.41 | Total tetracycline usage | 13366 |
| reg_f_acf1 | 0.93 | 0.61 | 1.07 | Use of automatic cluster flushing | 10392 |
| reg_f_calf_housing_type3 | 0.94 | 0.61 | 1.07 | Type of calf housing | 10640 |
| reg_weaned_heifer1 | 0.94 | 0.68 | 1.06 | Sample collected from the environment of a weaned heifer | 14146 |
| reg_weaned_heifer0 | 1.06 | 0.94 | 1.46 | Sample collected from the environment of a weaned heifer | 14252 |
| reg_f_poultry1 | 1.06 | 0.94 | 1.59 | Presence of poultry on the farm | 10856 |
| reg_f_foragetype_3yes | 1.06 | 0.93 | 1.64 | Maize silage used on farm | 11536 |
| main_cefalexin | 0.95 | 0.68 | 1.32 | Total cefalexin usage | 12769 |
| reg_f_firstmastitisOrbeninLA | 1.05 | 0.86 | 2.19 | First line choice of mastitis treatment is cloxacillin | 12774 |
| reg_f_lime1 | 1.05 | 0.93 | 1.53 | Use of lime in milking cow housing | 12295 |
| reg_f_bedmilk_1yes | 0.95 | 0.66 | 1.08 | Use of sand in milking cow bedding | 11987 |
| reg_f_firstmastitisUbrolexin | 0.96 | 0.58 | 1.11 | First line choice of mastitis treatment is cefalexin/ kanamycin | 13189 |
| reg_f_manure_1yes | 1.05 | 0.93 | 1.51 | Use of slurry lagoon | 10583 |
| reg_f_clostvaccY | 0.96 | 0.64 | 1.1 | Routine use of clostridial vaccination | 13062 |
| reg_f_salmvaccY | 0.96 | 0.58 | 1.13 | Routine use of Salmonella vaccination | 11197 |
| reg_f_predip1 | 1.04 | 0.93 | 1.44 | Whether predip is used when milking | 11247 |
| reg_f_fox2 | 0.96 | 0.71 | 1.08 | Perceived fox population on farm | 13343 |
| reg_f_equine1 | 1.04 | 0.92 | 1.42 | Presence of horses on the farm | 14623 |
| reg_f_badger2 | 1.04 | 0.9 | 1.55 | Perceived badger population on the farm | 12196 |
| reg_f_deer2 | 0.97 | 0.68 | 1.09 | Perceived amount of deer on farmland | 11514 |
| reg_s_calving_now1 | 1.03 | 0.93 | 1.38 | Sample collected during the calving season | 13902 |
| reg_f_manure_9yes | 1.03 | 0.87 | 1.68 | Use of other manure management | 12921 |
| reg_f_cleancluster.L | 0.97 | 0.76 | 1.07 | How often the milking cluster was cleaned | 13794 |
| reg_f_foragetype_8yes | 0.97 | 0.6 | 1.16 | Feeding of other forage | 14145 |
| reg_f_nsaidmast | 0.97 | 0.8 | 1.05 | Routine use of anti-inflammatories in cases of mastitis | 12903 |
| reg_f_shoot2 | 0.97 | 0.71 | 1.08 | Whether the shoot crosses the farm | 10858 |
| reg_f_firstmastitisMastiplanLC | 0.97 | 0.61 | 1.17 | First line choice of mastitis treatment is cefapirin | 15368 |
| reg_f_pattern2 | 1.03 | 0.91 | 1.43 | Calving pattern | 13418 |
| reg_f_whichclinrespPenicillimoxycillin | 0.97 | 0.62 | 1.17 | First line choice of pneumonia treatment is amoxicillin | 14627 |
| reg_f_machinery1 | 0.97 | 0.74 | 1.08 | Whether machinery was shared with other farms | 13809 |
| reg_f_anything_waste1 | 1.03 | 0.92 | 1.35 | Whether waste milk was fed to any calves | 14253 |
| reg_f_diarrvaccY | 0.97 | 0.74 | 1.09 | Routine use of calf enteric disease vaccination | 13367 |
| reg_f_percentdryoff | 1.03 | 0.95 | 1.22 | Proportion of cows dried off with antibiotic tubes | 13079 |
| reg_f_bedmilk_6yes | 1.03 | 0.86 | 1.54 | Use of green bedding for milking cows | 13859 |
| reg_f_manure_3yes | 1.03 | 0.91 | 1.38 | Use of a metal slurry store | 13836 |
| reg_f_othergrazeY | 0.97 | 0.75 | 1.08 | Use of grazing land away from the farm | 13721 |
| reg_f_rook2 | 1.03 | 0.91 | 1.34 | Perceived rook population on farm | 13452 |
| reg_f_feedlorry1 | 1.03 | 0.92 | 1.32 | Whether feed lorries crossed any cattle yards when delivering | 14284 |
| reg_f_geographical_area2 | 1.02 | 0.92 | 1.32 | Geographical area 2 | 13961 |
| reg_f_um_spray1 | 1.02 | 0.9 | 1.37 | Routine use of umbilical treatment at birth | 13855 |
| reg_f_organic1 | 0.98 | 0.71 | 1.13 | Whether the farm is organic | 13493 |
| reg_f_manure_6yes | 1.02 | 0.92 | 1.29 | Use of muck heap on concrete | 15221 |
| reg_f_daystreatmast | 1.02 | 0.94 | 1.21 | Number of days mastitis is treated for | 15025 |
| reg_adult1 | 1.02 | 0.92 | 1.25 | Sample collected from the milking cow environment | 15001 |
| reg_f_hostfarmwalk1 | 1.02 | 0.91 | 1.29 | Whether a farm walk has taken place on the farm in the last 2 years | 14356 |
| reg_f_bedmilk_2yes | 1.02 | 0.91 | 1.27 | Use of straw as milking cow bedding | 14695 |
| reg_f_trough_clean.L | 1.02 | 0.93 | 1.22 | Daily cleaning of water troughs in calf housing | 13884 |
| reg_f_timesfirstmast | 0.98 | 0.85 | 1.06 | How often clinical mastitis cases were treated with intramammary tubes | 14080 |
| reg_f_hectares | 0.98 | 0.85 | 1.06 | Size of farm in hectares | 13599 |
| reg_f_foragetype_4yes | 1.02 | 0.92 | 1.25 | Feeding of whole crop silage | 14484 |
| reg_f_time_dam.L | 1.02 | 0.92 | 1.24 | Time calf spends with dam | 13291 |
| reg_f_abcalve1 | 1.02 | 0.89 | 1.31 | Routine use of antibiotics following a difficult calving | 13603 |
| reg_f_manure_7yes | 1.02 | 0.86 | 1.37 | Use of composter for muck management | 15307 |
| reg_f_whichclinrespOtherDdontknow | 1.02 | 0.84 | 1.42 | First line choice of pneumonia treatment is unknown | 15451 |
| reg_s_rain | 1.02 | 0.96 | 1.12 | Average monthly rainfall | 15328 |
| reg_f_yield | 1.02 | 0.93 | 1.18 | 305 day milk yield | 14634 |
| reg_f_foragetype_1yes | 1.02 | 0.87 | 1.33 | Feeding of clamp grass silage | 15498 |
| reg_f_outsource1 | 1.01 | 0.88 | 1.3 | Whether heifer calves are reared by a different enterprise | 15180 |
| reg_f_firstmastitisCobactan | 0.99 | 0.73 | 1.19 | First line choice of mastitis treatment is cefquinome | 14799 |
| reg_f_aiexternal1 | 1.01 | 0.91 | 1.24 | Artificial insemination performed by external company | 14228 |
| reg_f_wean2 | 0.99 | 0.79 | 1.13 | Age of weaning | 14301 |
| reg_f_hunt2 | 0.99 | 0.81 | 1.1 | Whether the hunt crosses the farm | 14843 |
| reg_f_bedmilk_8yes | 0.99 | 0.72 | 1.21 | Unknown milking cow bedding type | 14857 |
| reg_f_halocur1 | 1.01 | 0.9 | 1.23 | Routine use of treatment to prevent cryptosporidiosis | 15015 |
| reg_f_muckspreader3 | 0.99 | 0.75 | 1.19 | Use of a contract muck spreader | 15745 |
| reg_f_bedmilk_4yes | 1.01 | 0.84 | 1.35 | Use of paper waste for milking cow bedding | 14530 |
| reg_f_calf_housing_older1 | 0.99 | 0.8 | 1.12 | Whether calves have been kept near to older animals | 15647 |
| reg_f_foragetype_7yes | 0.99 | 0.81 | 1.12 | Feeding of haylage | 14456 |
| reg_f_calf_housing_type2 | 1.01 | 0.9 | 1.21 | Type of calf housing | 15435 |
| reg_f_injectmast | 0.99 | 0.85 | 1.09 | Whether mastitis cases are routinely injected with antibiotic | 15577 |
| reg_f_nsaiddiarr1 | 0.99 | 0.82 | 1.12 | Routine use of anti-inflammatories in cases of calf diarrhoea | 14910 |
| reg_f_calving_group1 | 1.01 | 0.87 | 1.25 | Whether calves were born in a group pen | 14965 |
| reg_f_ibrvaccY | 0.99 | 0.82 | 1.11 | Routine use of IBR vaccination | 15763 |
| reg_dry1 | 0.99 | 0.83 | 1.11 | Sample collected from the dry cow environment | 15410 |
| reg_f_bought_pre | 1.01 | 0.93 | 1.14 | Number of cattle bought in the 12 months before the start of the project | 14805 |
| reg_f_foragetype_6yes | 1.01 | 0.87 | 1.26 | Feeding of straw | 15578 |
| reg_f_drytrough.L | 0.99 | 0.86 | 1.09 | How often the dry cow water trough was cleaned out | 14593 |
| reg_f_pneum_vacc1 | 1.01 | 0.89 | 1.21 | Routine use of calf respiratory vaccination | 15422 |
| reg_f_premises2 | 0.99 | 0.84 | 1.12 | Number of farm premises | 15055 |
| reg_f_market1 | 0.99 | 0.83 | 1.13 | Whether animals are taken to market | 14304 |
| reg_f_geographical_area3 | 1.01 | 0.88 | 1.19 | Geographical area 3 | 15533 |
| reg_f_bvdvaccY | 1.01 | 0.89 | 1.19 | Routine use of BVD vaccination | 14765 |
| reg_f_pheasant2 | 1.01 | 0.87 | 1.22 | Perceived pheasant population on farm | 16033 |
| reg_f_fram_dct1 | 0.99 | 0.84 | 1.13 | Use of framycetin dry cow therapy | 15346 |
| reg_f_firstmastitisUbroYellow | 0.99 | 0.82 | 1.14 | First line choice of mastitis treatment is streptomycin/ framycetin | 15943 |
| reg_f_rat2 | 1.01 | 0.89 | 1.19 | Perceived rat population on farm | 15668 |
| reg_f_wean3 | 0.99 | 0.85 | 1.12 | Age of weaning | 16139 |
| reg_footpath1 | 1.01 | 0.89 | 1.19 | Sample collected from a public footpath | 14807 |
| reg_f_herd_size | 1.01 | 0.92 | 1.13 | Total milking cattle on farm | 15719 |
| reg_f_manure_8yes | 1.01 | 0.86 | 1.23 | Use of separator for muck management | 15645 |
| reg_f_heifers_waste1 | 1.01 | 0.88 | 1.19 | Whether waste milk has been fed to heifers | 16652 |
| reg_f_anticocc1 | 0.99 | 0.84 | 1.13 | Routine use of anticoccidials | 14850 |
| reg_f_manure_2yes | 0.99 | 0.82 | 1.16 | Use of a weeping wall for slurry management | 15670 |
| reg_f_calveclean.L | 0.99 | 0.86 | 1.12 | How often the calving pens are cleaned out | 15365 |
| reg_f_pigeon2 | 1 | 0.89 | 1.16 | Perceived pigeon population on farm | 16287 |
| reg_f_total_cattle | 1 | 0.91 | 1.13 | Total number of cattle on farm | 15650 |
| reg_f_give_col1 | 1 | 0.87 | 1.19 | Administration of colostrum within six hours of life | 16275 |
| reg_f_starling2 | 1 | 0.85 | 1.14 | Perceived amount of starling infestation | 15575 |
| reg_f_whichclinrespTetracycline | 1 | 0.84 | 1.15 | First line choice of pneumonia treatment is tetracycline | 15426 |
| reg_f_clox_dct1 | 1 | 0.84 | 1.15 | Use of cloxacillin dry cow therapy | 14993 |
| reg_f_leptovaccY | 1 | 0.85 | 1.16 | Routine use of leptospirosis vaccination | 15273 |
| reg_f_milkgraze1 | 1 | 0.84 | 1.17 | Whether milking cows graze | 15249 |
| reg_f_manure_5yes | 1 | 0.85 | 1.16 | Muck heap in field used | 16334 |
| reg_f_water1 | 1 | 0.87 | 1.17 | Water supply to the farm | 16330 |
| reg_f_ceph_dct1 | 1 | 0.87 | 1.18 | Use of cephalonium dry cow therapy | 15314 |
| reg_f_nsaidcalve1 | 1 | 0.87 | 1.14 | Routine use of anti-inflammatories at calving | 15448 |
| reg_f_whichclinrespMacrolide | 1 | 0.86 | 1.14 | First line choice of pneumonia treatment is Macrolide | 15614 |
| reg_f_cefq_dct1 | 1 | 0.84 | 1.17 | Use of cefquinome dry cow therapy | 15436 |
| reg_f_bedmilk_3yes | 1 | 0.86 | 1.14 | Use of sawdust as milking cow bedding | 14504 |
| reg_f_muckspreader1 | 1 | 0.86 | 1.15 | Use of a contract muck spreader | 15654 |
| reg_f_treatfoot1 | 1 | 0.87 | 1.14 | Use of a footbath for lameness outbreaks | 15255 |
| reg_f_foragetype_5yes | 1 | 0.86 | 1.15 | Feeding of hay | 15969 |
| reg_f_scc | 1 | 0.91 | 1.09 | Somatic cell count | 15473 |

**Ciprofloxacin**

| **Variable** | **Odds ratio** | **Lower CI** | **Upper CI** | **Description** | **Effective sample size** |
| --- | --- | --- | --- | --- | --- |
| reg_preweaned_heifer1 | 4.13 | 2.79 | 6.46 | Sample collected from the environment of a pre-weaned heifer | 15693 |
| main_s_temp | 2.14 | 1.63 | 2.87 | Average monthly temperature | 12028 |
| reg_f_calf_housing_type3 | 0.69 | 0.25 | 1.07 | Type of calf housing | 4232 |
| main_fq | 1.43 | 1 | 2.06 | Total fluoroquinolone usage | 10768 |
| reg_f_firstmastitisUbroYellow | 0.73 | 0.23 | 1.09 | First line choice of mastitis treatment is streptomycin/ framycetin | 7563 |
| reg_weaned_heifer1 | 0.75 | 0.36 | 1.11 | Sample collected from the environment of a weaned heifer | 6799 |
| reg_weaned_heifer0 | 1.32 | 0.89 | 2.73 | Sample collected from the environment of a weaned heifer | 6662 |
| reg_f_manure_5yes | 0.78 | 0.34 | 1.07 | Muck heap in field used | 5482 |
| reg_f_nsaiddiarr1 | 0.83 | 0.33 | 1.1 | Routine use of anti-inflammatories in cases of calf diarrhoea | 6236 |
| reg_f_foragetype_8yes | 1.19 | 0.82 | 7.3 | Feeding of other forage | 7416 |
| reg_f_starling2 | 1.17 | 0.92 | 2.45 | Perceived amount of starling infestation | 6626 |
| reg_f_um_spray1 | 1.17 | 0.88 | 3.45 | Routine use of umbilical treatment at birth | 6640 |
| reg_f_badger2 | 0.86 | 0.33 | 1.14 | Perceived badger population on the farm | 6769 |
| reg_f_equine1 | 1.15 | 0.92 | 2.26 | Presence of horses on the farm | 7623 |
| main_novobiocin | 1.15 | 0.82 | 1.61 | Total usage of novobiocin | 12967 |
| reg_s_rain | 1.14 | 0.99 | 1.4 | Average monthly rainfall | 11234 |
| reg_f_foragetype_6yes | 0.89 | 0.37 | 1.15 | Feeding of straw | 8766 |
| reg_f_calf_housing_older1 | 0.91 | 0.47 | 1.12 | Whether calves have been kept near to older animals | 10059 |
| main_total_mg_pcu | 1.1 | 0.81 | 1.51 | Total antibiotic usage | 10506 |
| reg_f_lime1 | 0.92 | 0.48 | 1.14 | Use of lime in milking cow housing | 8800 |
| reg_f_geographical_area2 | 0.92 | 0.51 | 1.12 | Geographical area 2 | 10736 |
| reg_f_shows1 | 0.92 | 0.39 | 1.19 | Whether animals are taken to shows | 9575 |
| reg_f_pneum_vacc1 | 1.08 | 0.9 | 1.91 | Routine use of calf respiratory vaccination | 10497 |
| reg_f_nsaidcalve1 | 0.93 | 0.55 | 1.11 | Routine use of anti-inflammatories at calving | 11633 |
| reg_f_daystreatmast | 0.94 | 0.67 | 1.08 | Number of days mastitis is treated for | 7167 |
| main_ceph_3_4 | 1.07 | 0.79 | 1.44 | Total usage of third and fourth generation cephalosporins | 13594 |
| reg_f_timesfirstmast | 1.07 | 0.94 | 1.42 | How often clinical mastitis cases were treated with intramammary tubes | 11138 |
| reg_f_organic1 | 1.07 | 0.82 | 2.29 | Whether the farm is organic | 12424 |
| reg_f_shoot2 | 1.06 | 0.88 | 1.78 | Whether the shoot crosses the farm | 9954 |
| reg_f_poultry1 | 0.94 | 0.58 | 1.13 | Presence of poultry on the farm | 10396 |
| reg_f_bedmilk_6yes | 1.06 | 0.79 | 2.28 | Use of green bedding for milking cows | 12193 |
| reg_f_geographical_area3 | 1.06 | 0.88 | 1.69 | Geographical area 3 | 11375 |
| reg_f_deer2 | 1.06 | 0.87 | 1.69 | Perceived amount of deer on farmland | 12957 |
| reg_f_fox2 | 1.05 | 0.86 | 1.76 | Perceived fox population on farm | 7675 |
| reg_f_foragetype_3yes | 0.95 | 0.58 | 1.15 | Maize silage used on farm | 10977 |
| reg_f_whichclinrespOtherDdontknow | 0.95 | 0.46 | 1.37 | First line choice of pneumonia treatment is unknown | 13574 |
| reg_f_manure_7yes | 1.05 | 0.81 | 1.93 | Use of composter for muck management | 13782 |
| reg_f_foragetype_4yes | 1.05 | 0.88 | 1.55 | Feeding of whole crop silage | 13965 |
| reg_f_manure_9yes | 0.96 | 0.49 | 1.28 | Use of other manure management | 13485 |
| reg_f_foragetype_7yes | 1.04 | 0.85 | 1.66 | Feeding of haylage | 11290 |
| reg_f_bedmilk_2yes | 0.96 | 0.62 | 1.16 | Use of straw as milking cow bedding | 13291 |
| reg_f_bedmilk_4yes | 1.04 | 0.78 | 1.96 | Use of paper waste for milking cow bedding | 13598 |
| reg_f_calf_housing_type2 | 1.04 | 0.84 | 1.63 | Type of calf housing | 12407 |
| reg_f_bedmilk_3yes | 0.96 | 0.65 | 1.16 | Use of sawdust as milking cow bedding | 13805 |
| reg_f_clox_dct1 | 1.04 | 0.86 | 1.55 | Use of cloxacillin dry cow therapy | 14131 |
| reg_f_market1 | 0.96 | 0.66 | 1.15 | Whether animals are taken to market | 12646 |
| reg_f_hectares | 0.96 | 0.75 | 1.08 | Size of farm in hectares | 13697 |
| reg_f_wean2 | 1.04 | 0.83 | 1.61 | Age of weaning | 14120 |
| reg_adult1 | 1.04 | 0.86 | 1.48 | Sample collected from the milking cow environment | 13968 |
| reg_f_machinery1 | 1.04 | 0.87 | 1.48 | Whether machinery was shared with other farms | 14737 |
| reg_f_manure_6yes | 0.97 | 0.66 | 1.17 | Use of muck heap on concrete | 10417 |
| reg_f_nsaidmast | 0.97 | 0.77 | 1.1 | Routine use of anti-inflammatories in cases of mastitis | 14039 |
| reg_f_time_dam.L | 1.03 | 0.89 | 1.4 | Time calf spends with dam | 13189 |
| reg_f_whichclinrespTetracycline | 1.03 | 0.84 | 1.5 | First line choice of pneumonia treatment is tetracycline | 14007 |
| reg_f_predip1 | 1.03 | 0.85 | 1.48 | Whether predip is used when milking | 14150 |
| reg_f_anything_waste1 | 1.03 | 0.85 | 1.47 | Whether waste milk was fed to any calves | 13999 |
| reg_f_rat2 | 1.03 | 0.86 | 1.45 | Perceived rat population on farm | 12527 |
| reg_f_drytrough.L | 1.03 | 0.88 | 1.35 | How often the dry cow water trough was cleaned out | 13553 |
| reg_f_othergrazeY | 1.03 | 0.85 | 1.45 | Use of grazing land away from the farm | 14048 |
| reg_f_firstmastitistdDmultiject | 1.03 | 0.84 | 1.49 | First line choice of mastitis treatment is streptomycin/ neomycin/ novobiocin/ penicillin | 13907 |
| reg_f_salmvaccY | 0.97 | 0.6 | 1.31 | Routine use of Salmonella vaccination | 14613 |
| reg_s_housed_outdooroutdoor | 0.97 | 0.74 | 1.14 | Sample collected from pasture | 15477 |
| reg_f_bedmilk_1yes | 1.03 | 0.84 | 1.47 | Use of sand in milking cow bedding | 14998 |
| reg_f_firstmastitisOrbeninLA | 0.98 | 0.57 | 1.38 | First line choice of mastitis treatment is cloxacillin | 14319 |
| reg_f_whichclinrespMacrolide | 1.02 | 0.85 | 1.42 | First line choice of pneumonia treatment is Macrolide | 14843 |
| reg_f_foragetype_1yes | 1.02 | 0.77 | 1.6 | Feeding of clamp grass silage | 14457 |
| reg_f_bought_pre | 0.98 | 0.79 | 1.11 | Number of cattle bought in the 12 months before the start of the project | 14481 |
| reg_f_yield | 1.02 | 0.9 | 1.27 | 305 day milk yield | 13160 |
| reg_f_muckspreader1 | 0.98 | 0.72 | 1.18 | Use of a contract muck spreader | 14627 |
| reg_f_calveclean.L | 1.02 | 0.86 | 1.35 | How often the calving pens are cleaned out | 14038 |
| reg_f_halocur1 | 0.98 | 0.72 | 1.18 | Routine use of treatment to prevent cryptosporidiosis | 15356 |
| reg_f_wean3 | 1.02 | 0.83 | 1.42 | Age of weaning | 13750 |
| reg_f_manure_8yes | 0.98 | 0.66 | 1.26 | Use of separator for muck management | 15110 |
| reg_f_manure_2yes | 0.98 | 0.68 | 1.23 | Use of a weeping wall for slurry management | 13636 |
| reg_f_cleancluster.L | 0.98 | 0.77 | 1.14 | How often the milking cluster was cleaned | 15715 |
| reg_f_firstmastitisCobactan | 1.02 | 0.75 | 1.61 | First line choice of mastitis treatment is cefquinome | 14490 |
| reg_f_firstmastitisUbrolexin | 0.98 | 0.66 | 1.27 | First line choice of mastitis treatment is cefalexin/ kanamycin | 15576 |
| reg_f_injectmast | 0.98 | 0.79 | 1.13 | Whether mastitis cases are routinely injected with antibiotic | 14950 |
| reg_f_premises2 | 0.98 | 0.72 | 1.19 | Number of farm premises | 13996 |
| reg_f_manure_1yes | 1.02 | 0.84 | 1.38 | Use of slurry lagoon | 15333 |
| reg_s_calving_now1 | 1.02 | 0.85 | 1.36 | Sample collected during the calving season | 15832 |
| reg_f_scc | 0.98 | 0.79 | 1.12 | Somatic cell count | 13187 |
| reg_f_manure_3yes | 0.98 | 0.71 | 1.21 | Use of a metal slurry store | 14400 |
| reg_f_hunt2 | 0.98 | 0.71 | 1.21 | Whether the hunt crosses the farm | 15981 |
| reg_f_rook2 | 1.02 | 0.82 | 1.41 | Perceived rook population on farm | 15432 |
| reg_f_percentdryoff | 0.98 | 0.82 | 1.11 | Proportion of cows dried off with antibiotic tubes | 13978 |
| reg_footpath1 | 1.02 | 0.85 | 1.32 | Sample collected from a public footpath | 15593 |
| reg_f_hostfarmwalk1 | 1.02 | 0.83 | 1.36 | Whether a farm walk has taken place on the farm in the last 2 years | 15339 |
| reg_f_clostvaccY | 0.98 | 0.71 | 1.23 | Routine use of clostridial vaccination | 15736 |
| reg_f_manure_4yes | 0.98 | 0.76 | 1.18 | Use of a concrete slurry store | 15247 |
| reg_f_heifers_waste1 | 1.02 | 0.82 | 1.37 | Whether waste milk has been fed to heifers | 15220 |
| reg_f_foragetype_2yes | 1.01 | 0.8 | 1.41 | Feeding of big bale grass silage | 15358 |
| reg_f_treatfoot1 | 0.99 | 0.76 | 1.21 | Use of a footbath for lameness outbreaks | 15499 |
| reg_f_pheasant2 | 0.99 | 0.73 | 1.25 | Perceived pheasant population on farm | 15243 |
| reg_f_lungvaccY | 1.01 | 0.83 | 1.32 | Routine use of lungworm vaccination | 15349 |
| reg_f_feedlorry1 | 0.99 | 0.76 | 1.22 | Whether feed lorries crossed any cattle yards when delivering | 15026 |
| reg_f_leptovaccY | 1.01 | 0.81 | 1.36 | Routine use of leptospirosis vaccination | 14451 |
| reg_f_calving_group1 | 0.99 | 0.7 | 1.29 | Whether calves were born in a group pen | 15731 |
| reg_f_bedmilk_8yes | 1.01 | 0.69 | 1.58 | Unknown milking cow bedding type | 16431 |
| reg_f_outsource1 | 0.99 | 0.72 | 1.28 | Whether heifer calves are reared by a different enterprise | 16010 |
| reg_f_total_cattle | 0.99 | 0.82 | 1.16 | Total number of cattle on farm | 14539 |
| reg_f_abcalve1 | 1.01 | 0.78 | 1.37 | Routine use of antibiotics following a difficult calving | 14795 |
| reg_f_diarrvaccY | 0.99 | 0.77 | 1.21 | Routine use of calf enteric disease vaccination | 16283 |
| reg_f_anticocc1 | 0.99 | 0.75 | 1.23 | Routine use of anticoccidials | 14699 |
| reg_f_ceph_dct1 | 1.01 | 0.81 | 1.33 | Use of cephalonium dry cow therapy | 14106 |
| reg_f_give_col1 | 0.99 | 0.76 | 1.24 | Administration of colostrum within six hours of life | 15443 |
| reg_f_pigeon2 | 1.01 | 0.82 | 1.28 | Perceived pigeon population on farm | 15848 |
| reg_f_trough_clean.L | 1.01 | 0.85 | 1.25 | Daily cleaning of water troughs in calf housing | 14505 |
| reg_f_herd_size | 1.01 | 0.86 | 1.22 | Total milking cattle on farm | 15973 |
| reg_dry1 | 0.99 | 0.77 | 1.24 | Sample collected from the dry cow environment | 16680 |
| reg_f_acf1 | 1.01 | 0.81 | 1.31 | Use of automatic cluster flushing | 16258 |
| reg_f_cefq_dct1 | 0.99 | 0.75 | 1.26 | Use of cefquinome dry cow therapy | 15543 |
| reg_f_pattern2 | 1.01 | 0.8 | 1.32 | Calving pattern | 16463 |
| reg_f_milkgraze1 | 1.01 | 0.77 | 1.38 | Whether milking cows graze | 15393 |
| reg_f_whichclinrespPenicillimoxycillin | 0.99 | 0.64 | 1.45 | First line choice of pneumonia treatment is amoxicillin | 16502 |
| reg_f_firstmastitisMastiplanLC | 0.99 | 0.64 | 1.45 | First line choice of mastitis treatment is cefapirin | 16583 |
| reg_f_bvdvaccY | 1.01 | 0.81 | 1.29 | Routine use of BVD vaccination | 15891 |
| reg_f_water1 | 1 | 0.81 | 1.28 | Water supply to the farm | 16726 |
| reg_f_ibrvaccY | 1 | 0.79 | 1.22 | Routine use of IBR vaccination | 16969 |
| reg_f_foragetype_5yes | 1 | 0.79 | 1.32 | Feeding of hay | 16330 |
| reg_f_muckspreader3 | 1 | 0.65 | 1.49 | Use of a contract muck spreader | 15633 |
| reg_f_aiexternal1 | 1 | 0.79 | 1.23 | Artificial insemination performed by external company | 15194 |
| reg_f_fram_dct1 | 1 | 0.79 | 1.27 | Use of framycetin dry cow therapy | 16268 |

**Streptomycin**

| **Variable** | **Odds ratio** | **Lower CI** | **Upper CI** | **Description** | **Effective sample size** |
| --- | --- | --- | --- | --- | --- |
| reg_preweaned_heifer1 | 1.95 | 1.46 | 2.51 | Sample collected from the environment of a pre-weaned heifer | 15464 |
| reg_footpath1 | 0.57 | 0.33 | 1.01 | Sample collected from a public footpath | 8540 |
| main_s_temp | 1.53 | 1.32 | 1.77 | Average monthly temperature | 13407 |
| reg_s_housed_outdooroutdoor | 0.75 | 0.44 | 1.03 | Sample collected from pasture | 8326 |
| reg_s_calving_now1 | 1.27 | 0.98 | 2 | Sample collected during the calving season | 8738 |
| reg_dry1 | 1.16 | 0.97 | 1.79 | Sample collected from the dry cow environment | 12817 |
| reg_f_foragetype_3yes | 1.14 | 0.95 | 2.08 | Maize silage used on farm | 5522 |
| reg_f_pneum_vacc1 | 1.14 | 0.95 | 1.92 | Routine use of calf respiratory vaccination | 6699 |
| main_total_mg_pcu | 1.13 | 0.86 | 1.49 | Total antibiotic usage | 12578 |
| reg_f_abcalve1 | 0.89 | 0.48 | 1.07 | Routine use of antibiotics following a difficult calving | 10026 |
| main_cefalexin | 0.9 | 0.75 | 1.09 | Total cefalexin usage | 12778 |
| reg_f_wean2 | 0.9 | 0.52 | 1.06 | Age of weaning | 10638 |
| reg_f_firstmastitisUbrolexin | 0.91 | 0.44 | 1.09 | First line choice of mastitis treatment is cefalexin/ kanamycin | 10314 |
| main_tet | 1.09 | 0.89 | 1.34 | Total tetracycline usage | 12151 |
| reg_f_cleancluster.L | 0.92 | 0.69 | 1.04 | How often the milking cluster was cleaned | 11173 |
| reg_f_foragetype_5yes | 1.08 | 0.94 | 1.65 | Feeding of hay | 11173 |
| reg_f_bedmilk_4yes | 1.07 | 0.89 | 2.5 | Use of paper waste for milking cow bedding | 9850 |
| reg_f_pheasant2 | 1.06 | 0.93 | 1.69 | Perceived pheasant population on farm | 10567 |
| reg_f_shows1 | 1.06 | 0.91 | 1.79 | Whether animals are taken to shows | 12426 |
| reg_f_halocur1 | 0.95 | 0.66 | 1.07 | Routine use of treatment to prevent cryptosporidiosis | 10267 |
| reg_f_nsaidmast | 0.95 | 0.77 | 1.04 | Routine use of anti-inflammatories in cases of mastitis | 12851 |
| reg_adult1 | 0.95 | 0.73 | 1.04 | Sample collected from the milking cow environment | 13692 |
| reg_f_whichclinrespPenicillimoxycillin | 0.95 | 0.47 | 1.16 | First line choice of pneumonia treatment is amoxicillin | 12856 |
| reg_f_firstmastitisMastiplanLC | 0.95 | 0.48 | 1.15 | First line choice of mastitis treatment is cefapirin | 12121 |
| reg_f_organic1 | 0.96 | 0.59 | 1.13 | Whether the farm is organic | 11453 |
| reg_f_firstmastitistdDmultiject | 1.04 | 0.93 | 1.44 | First line choice of mastitis treatment is streptomycin/ neomycin/ novobiocin/ penicillin | 12760 |
| reg_f_ceph_dct1 | 1.04 | 0.93 | 1.41 | Use of cephalonium dry cow therapy | 13189 |
| reg_f_manure_3yes | 1.04 | 0.92 | 1.44 | Use of a metal slurry store | 10961 |
| reg_f_milkgraze1 | 1.04 | 0.91 | 1.5 | Whether milking cows graze | 10565 |
| reg_f_foragetype_2yes | 0.96 | 0.67 | 1.1 | Feeding of big bale grass silage | 12546 |
| reg_f_total_cattle | 1.04 | 0.96 | 1.26 | Total number of cattle on farm | 12890 |
| reg_f_foragetype_8yes | 0.97 | 0.6 | 1.16 | Feeding of other forage | 13344 |
| reg_f_geographical_area2 | 0.97 | 0.74 | 1.08 | Geographical area 2 | 13343 |
| reg_f_whichclinrespTetracycline | 0.97 | 0.72 | 1.08 | First line choice of pneumonia treatment is tetracycline | 13601 |
| main_amox | 1.03 | 0.84 | 1.28 | Total amoxicillin usage | 12754 |
| reg_f_cefq_dct1 | 0.97 | 0.72 | 1.09 | Use of cefquinome dry cow therapy | 13947 |
| reg_f_bvdvaccY | 1.03 | 0.93 | 1.35 | Routine use of BVD vaccination | 12468 |
| reg_f_time_dam.L | 1.03 | 0.93 | 1.34 | Time calf spends with dam | 8561 |
| reg_f_leptovaccY | 1.03 | 0.92 | 1.38 | Routine use of leptospirosis vaccination | 11081 |
| reg_f_heifers_waste1 | 1.03 | 0.92 | 1.36 | Whether waste milk has been fed to heifers | 10976 |
| reg_f_rook2 | 1.03 | 0.92 | 1.32 | Perceived rook population on farm | 14780 |
| reg_f_muckspreader3 | 0.97 | 0.65 | 1.16 | Use of a contract muck spreader | 14810 |
| reg_f_foragetype_6yes | 1.03 | 0.89 | 1.43 | Feeding of straw | 13953 |
| reg_f_acf1 | 0.98 | 0.76 | 1.1 | Use of automatic cluster flushing | 14077 |
| reg_f_calf_housing_type2 | 0.98 | 0.79 | 1.08 | Type of calf housing | 14890 |
| reg_f_aiexternal1 | 1.02 | 0.93 | 1.28 | Artificial insemination performed by external company | 14294 |
| reg_f_fram_dct1 | 0.98 | 0.76 | 1.09 | Use of framycetin dry cow therapy | 12804 |
| reg_f_calveclean.L | 1.02 | 0.93 | 1.27 | How often the calving pens are cleaned out | 14198 |
| reg_f_othergrazeY | 0.98 | 0.78 | 1.09 | Use of grazing land away from the farm | 14255 |
| reg_f_herd_size | 1.02 | 0.94 | 1.2 | Total milking cattle on farm | 15398 |
| reg_f_foragetype_1yes | 1.02 | 0.88 | 1.37 | Feeding of clamp grass silage | 14665 |
| reg_f_scc | 0.98 | 0.85 | 1.05 | Somatic cell count | 15049 |
| reg_f_clostvaccY | 1.02 | 0.9 | 1.27 | Routine use of clostridial vaccination | 14789 |
| reg_f_trough_clean.L | 1.02 | 0.93 | 1.2 | Daily cleaning of water troughs in calf housing | 14231 |
| reg_f_starling2 | 1.02 | 0.91 | 1.24 | Perceived amount of starling infestation | 14589 |
| reg_f_equine1 | 0.98 | 0.81 | 1.1 | Presence of horses on the farm | 13757 |
| reg_f_whichclinrespMacrolide | 0.98 | 0.82 | 1.09 | First line choice of pneumonia treatment is Macrolide | 14467 |
| reg_f_calving_group1 | 1.02 | 0.89 | 1.29 | Whether calves were born in a group pen | 14737 |
| reg_f_foragetype_4yes | 0.99 | 0.83 | 1.09 | Feeding of whole crop silage | 14526 |
| reg_f_daystreatmast | 1.01 | 0.94 | 1.15 | Number of days mastitis is treated for | 14779 |
| reg_f_machinery1 | 1.01 | 0.91 | 1.22 | Whether machinery was shared with other farms | 15002 |
| reg_f_percentdryoff | 1.01 | 0.95 | 1.14 | Proportion of cows dried off with antibiotic tubes | 15287 |
| reg_f_manure_1yes | 0.99 | 0.82 | 1.1 | Use of slurry lagoon | 12786 |
| reg_f_feedlorry1 | 0.99 | 0.82 | 1.1 | Whether feed lorries crossed any cattle yards when delivering | 15377 |
| reg_f_manure_6yes | 1.01 | 0.9 | 1.22 | Use of muck heap on concrete | 13867 |
| reg_f_bedmilk_8yes | 0.99 | 0.74 | 1.21 | Unknown milking cow bedding type | 14951 |
| reg_f_pigeon2 | 1.01 | 0.91 | 1.19 | Perceived pigeon population on farm | 15008 |
| reg_f_manure_4yes | 1.01 | 0.91 | 1.19 | Use of a concrete slurry store | 15383 |
| reg_f_deer2 | 1.01 | 0.9 | 1.22 | Perceived amount of deer on farmland | 15014 |
| main_strep | 1.01 | 0.84 | 1.22 | Total streptomycin usage | 12830 |
| reg_f_predip1 | 1.01 | 0.92 | 1.19 | Whether predip is used when milking | 15077 |
| reg_f_pattern2 | 0.99 | 0.81 | 1.12 | Calving pattern | 13256 |
| reg_f_geographical_area3 | 1.01 | 0.9 | 1.21 | Geographical area 3 | 14418 |
| reg_f_um_spray1 | 1.01 | 0.88 | 1.23 | Routine use of umbilical treatment at birth | 15700 |
| reg_f_salmvaccY | 1.01 | 0.85 | 1.31 | Routine use of Salmonella vaccination | 14891 |
| reg_f_bedmilk_1yes | 0.99 | 0.83 | 1.11 | Use of sand in milking cow bedding | 14669 |
| reg_f_lungvaccY | 1.01 | 0.9 | 1.19 | Routine use of lungworm vaccination | 14451 |
| reg_weaned_heifer1 | 1.01 | 0.9 | 1.18 | Sample collected from the environment of a weaned heifer | 14119 |
| reg_weaned_heifer0 | 0.99 | 0.85 | 1.1 | Sample collected from the environment of a weaned heifer | 14629 |
| reg_f_yield | 1.01 | 0.93 | 1.14 | 305 day milk yield | 14481 |
| reg_f_premises2 | 0.99 | 0.84 | 1.11 | Number of farm premises | 14509 |
| reg_f_badger2 | 0.99 | 0.81 | 1.14 | Perceived badger population on the farm | 14838 |
| reg_f_anticocc1 | 1.01 | 0.89 | 1.2 | Routine use of anticoccidials | 15149 |
| reg_f_bought_pre | 1.01 | 0.93 | 1.13 | Number of cattle bought in the 12 months before the start of the project | 16151 |
| reg_s_rain | 1.01 | 0.97 | 1.08 | Average monthly rainfall | 15991 |
| reg_f_ibrvaccY | 1.01 | 0.9 | 1.19 | Routine use of IBR vaccination | 15637 |
| reg_f_firstmastitisCobactan | 0.99 | 0.78 | 1.19 | First line choice of mastitis treatment is cefquinome | 14987 |
| reg_f_nsaidcalve1 | 1.01 | 0.9 | 1.18 | Routine use of anti-inflammatories at calving | 14262 |
| reg_f_rat2 | 1.01 | 0.9 | 1.18 | Perceived rat population on farm | 14971 |
| reg_f_diarrvaccY | 1.01 | 0.9 | 1.18 | Routine use of calf enteric disease vaccination | 15340 |
| reg_f_nsaiddiarr1 | 1.01 | 0.89 | 1.19 | Routine use of anti-inflammatories in cases of calf diarrhoea | 15850 |
| reg_f_manure_2yes | 1.01 | 0.88 | 1.21 | Use of a weeping wall for slurry management | 14431 |
| reg_f_drytrough.L | 0.99 | 0.87 | 1.09 | How often the dry cow water trough was cleaned out | 14578 |
| reg_f_injectmast | 0.99 | 0.87 | 1.09 | Whether mastitis cases are routinely injected with antibiotic | 14922 |
| reg_f_wean3 | 0.99 | 0.85 | 1.12 | Age of weaning | 15060 |
| reg_f_give_col1 | 1.01 | 0.89 | 1.18 | Administration of colostrum within six hours of life | 14275 |
| reg_f_shoot2 | 1.01 | 0.88 | 1.19 | Whether the shoot crosses the farm | 14900 |
| reg_f_bedmilk_2yes | 1.01 | 0.89 | 1.18 | Use of straw as milking cow bedding | 15338 |
| reg_f_treatfoot1 | 1.01 | 0.9 | 1.17 | Use of a footbath for lameness outbreaks | 14794 |
| reg_f_manure_8yes | 1.01 | 0.86 | 1.23 | Use of separator for muck management | 16202 |
| reg_f_whichclinrespOtherDdontknow | 0.99 | 0.77 | 1.22 | First line choice of pneumonia treatment is unknown | 16004 |
| reg_f_manure_9yes | 1.01 | 0.84 | 1.26 | Use of other manure management | 15507 |
| reg_f_firstmastitisOrbeninLA | 1.01 | 0.81 | 1.3 | First line choice of mastitis treatment is cloxacillin | 15499 |
| reg_f_muckspreader1 | 1.01 | 0.89 | 1.17 | Use of a contract muck spreader | 15809 |
| reg_f_foragetype_7yes | 1.01 | 0.88 | 1.18 | Feeding of haylage | 15916 |
| reg_f_poultry1 | 1.01 | 0.89 | 1.18 | Presence of poultry on the farm | 13463 |
| main_fq | 1 | 0.81 | 1.24 | Total fluoroquinolone usage | 12370 |
| reg_f_hectares | 1 | 0.93 | 1.1 | Size of farm in hectares | 15000 |
| reg_f_anything_waste1 | 1 | 0.86 | 1.13 | Whether waste milk was fed to any calves | 16071 |
| reg_f_manure_7yes | 1 | 0.83 | 1.25 | Use of composter for muck management | 15095 |
| reg_f_bedmilk_3yes | 1 | 0.87 | 1.12 | Use of sawdust as milking cow bedding | 15988 |
| reg_f_timesfirstmast | 1 | 0.91 | 1.08 | How often clinical mastitis cases were treated with intramammary tubes | 15557 |
| reg_f_manure_5yes | 1 | 0.86 | 1.13 | Muck heap in field used | 16106 |
| reg_f_outsource1 | 1 | 0.83 | 1.17 | Whether heifer calves are reared by a different enterprise | 15346 |
| reg_f_hunt2 | 1 | 0.89 | 1.15 | Whether the hunt crosses the farm | 15077 |
| reg_f_calf_housing_type3 | 1 | 0.86 | 1.13 | Type of calf housing | 16222 |
| reg_f_hostfarmwalk1 | 1 | 0.88 | 1.17 | Whether a farm walk has taken place on the farm in the last 2 years | 15878 |
| reg_f_water1 | 1 | 0.85 | 1.14 | Water supply to the farm | 15786 |
| reg_f_clox_dct1 | 1 | 0.86 | 1.15 | Use of cloxacillin dry cow therapy | 14865 |
| reg_f_lime1 | 1 | 0.88 | 1.15 | Use of lime in milking cow housing | 15821 |
| reg_f_firstmastitisUbroYellow | 1 | 0.87 | 1.17 | First line choice of mastitis treatment is streptomycin/ framycetin | 14361 |
| reg_f_fox2 | 1 | 0.87 | 1.14 | Perceived fox population on farm | 15965 |
| reg_f_calf_housing_older1 | 1 | 0.87 | 1.16 | Whether calves have been kept near to older animals | 14803 |
| reg_f_market1 | 1 | 0.87 | 1.15 | Whether animals are taken to market | 16430 |
| reg_f_bedmilk_6yes | 1 | 0.8 | 1.26 | Use of green bedding for milking cows | 15380 |

**Tetracycline**

| **Variable** | **Odds ratio** | **Lower CI** | **Upper CI** | **Description** | **Effective sample size** |
| --- | --- | --- | --- | --- | --- |
| reg_s_housed_outdooroutdoor | 0.24 | 0.15 | 0.35 | Sample collected from pasture | 14783 |
| reg_preweaned_heifer1 | 3.39 | 2 | 5.82 | Sample collected from the environment of a pre-weaned heifer | 16320 |
| reg_f_foragetype_3yes | 2.15 | 0.98 | 4.53 | Maize silage used on farm | 5799 |
| reg_s_calving_now1 | 2.02 | 1.1 | 3.29 | Sample collected during the calving season | 11454 |
| main_s_temp | 1.98 | 1.55 | 2.55 | Average monthly temperature | 11857 |
| reg_f_foragetype_2yes | 0.67 | 0.22 | 1.09 | Feeding of big bale grass silage | 7025 |
| main_total_mg_pcu | 1.47 | 1.01 | 2.13 | Total antibiotic usage | 10534 |
| reg_f_calf_housing_older1 | 0.74 | 0.38 | 1.06 | Whether calves have been kept near to older animals | 9125 |
| reg_dry1 | 0.74 | 0.42 | 1.05 | Sample collected from the dry cow environment | 12095 |
| main_strep | 0.76 | 0.6 | 0.97 | Total streptomycin usage | 10355 |
| reg_s_rain | 1.26 | 1.08 | 1.47 | Average monthly rainfall | 15742 |
| reg_weaned_heifer1 | 1.26 | 0.87 | 2.27 | Sample collected from the environment of a weaned heifer | 9246 |
| reg_weaned_heifer0 | 0.8 | 0.44 | 1.15 | Sample collected from the environment of a weaned heifer | 9151 |
| main_amox | 0.83 | 0.6 | 1.15 | Total amoxicillin usage | 10154 |
| reg_f_calf_housing_type3 | 0.84 | 0.43 | 1.1 | Type of calf housing | 10060 |
| reg_f_manure_7yes | 1.18 | 0.83 | 3.86 | Use of composter for muck management | 10262 |
| reg_f_lime1 | 1.18 | 0.92 | 2.14 | Use of lime in milking cow housing | 9542 |
| reg_f_yield | 1.17 | 0.96 | 1.76 | 305 day milk yield | 6215 |
| reg_footpath1 | 1.15 | 0.91 | 2.02 | Sample collected from a public footpath | 13684 |
| reg_f_geographical_area2 | 1.14 | 0.91 | 2.05 | Geographical area 2 | 10536 |
| reg_f_calf_housing_type2 | 1.13 | 0.9 | 1.93 | Type of calf housing | 10850 |
| reg_f_clostvaccY | 1.12 | 0.88 | 2.05 | Routine use of clostridial vaccination | 12058 |
| reg_f_firstmastitistdDmultiject | 1.11 | 0.89 | 1.86 | First line choice of mastitis treatment is streptomycin/ neomycin/ novobiocin/ penicillin | 11310 |
| reg_f_shows1 | 1.1 | 0.84 | 2.21 | Whether animals are taken to shows | 12045 |
| reg_f_bedmilk_8yes | 0.91 | 0.33 | 1.29 | Unknown milking cow bedding type | 12577 |
| reg_f_equine1 | 1.1 | 0.9 | 1.74 | Presence of horses on the farm | 11693 |
| reg_f_firstmastitisUbrolexin | 0.92 | 0.45 | 1.19 | First line choice of mastitis treatment is cefalexin/ kanamycin | 11084 |
| reg_f_bedmilk_2yes | 1.09 | 0.89 | 1.82 | Use of straw as milking cow bedding | 12136 |
| reg_f_foragetype_7yes | 0.92 | 0.55 | 1.14 | Feeding of haylage | 13396 |
| reg_f_salmvaccY | 1.08 | 0.78 | 2.46 | Routine use of Salmonella vaccination | 11949 |
| reg_f_manure_4yes | 0.92 | 0.62 | 1.12 | Use of a concrete slurry store | 12254 |
| reg_f_whichclinrespPenicillimoxycillin | 0.92 | 0.35 | 1.31 | First line choice of pneumonia treatment is amoxicillin | 13006 |
| reg_f_firstmastitisMastiplanLC | 0.92 | 0.37 | 1.34 | First line choice of mastitis treatment is cefapirin | 11972 |
| reg_f_treatfoot1 | 1.07 | 0.88 | 1.65 | Use of a footbath for lameness outbreaks | 12678 |
| reg_f_percentdryoff | 1.07 | 0.95 | 1.34 | Proportion of cows dried off with antibiotic tubes | 13056 |
| reg_f_ceph_dct1 | 0.93 | 0.61 | 1.13 | Use of cephalonium dry cow therapy | 11352 |
| reg_f_foragetype_5yes | 1.07 | 0.88 | 1.66 | Feeding of hay | 13979 |
| reg_f_water1 | 1.07 | 0.87 | 1.67 | Water supply to the farm | 12733 |
| reg_f_hunt2 | 1.07 | 0.88 | 1.61 | Whether the hunt crosses the farm | 13310 |
| reg_f_cefq_dct1 | 1.06 | 0.85 | 1.72 | Use of cefquinome dry cow therapy | 13318 |
| reg_f_hectares | 1.06 | 0.94 | 1.33 | Size of farm in hectares | 12996 |
| reg_f_firstmastitisCobactan | 0.94 | 0.47 | 1.3 | First line choice of mastitis treatment is cefquinome | 14092 |
| reg_f_give_col1 | 1.06 | 0.86 | 1.62 | Administration of colostrum within six hours of life | 13436 |
| reg_f_foragetype_8yes | 0.95 | 0.47 | 1.33 | Feeding of other forage | 14021 |
| reg_f_pattern2 | 0.95 | 0.59 | 1.19 | Calving pattern | 13079 |
| reg_f_wean3 | 0.95 | 0.66 | 1.15 | Age of weaning | 13309 |
| reg_f_bought_pre | 0.95 | 0.76 | 1.09 | Number of cattle bought in the 12 months before the start of the project | 13511 |
| reg_f_machinery1 | 1.05 | 0.86 | 1.51 | Whether machinery was shared with other farms | 13494 |
| reg_f_anything_waste1 | 0.95 | 0.66 | 1.16 | Whether waste milk was fed to any calves | 13856 |
| reg_f_injectmast | 1.05 | 0.9 | 1.38 | Whether mastitis cases are routinely injected with antibiotic | 13948 |
| reg_f_foragetype_4yes | 1.05 | 0.87 | 1.45 | Feeding of whole crop silage | 14290 |
| main_cefalexin | 1.05 | 0.75 | 1.46 | Total cefalexin usage | 9197 |
| reg_f_poultry1 | 1.05 | 0.86 | 1.51 | Presence of poultry on the farm | 13159 |
| reg_f_muckspreader3 | 0.96 | 0.5 | 1.39 | Use of a contract muck spreader | 14964 |
| reg_f_cleancluster.L | 0.96 | 0.74 | 1.12 | How often the milking cluster was cleaned | 14549 |
| reg_f_ibrvaccY | 1.04 | 0.85 | 1.49 | Routine use of IBR vaccination | 13355 |
| reg_f_time_dam.L | 1.04 | 0.84 | 1.46 | Time calf spends with dam | 11342 |
| reg_f_anticocc1 | 1.04 | 0.84 | 1.51 | Routine use of anticoccidials | 14568 |
| reg_f_predip1 | 1.04 | 0.86 | 1.42 | Whether predip is used when milking | 14122 |
| reg_f_shoot2 | 0.96 | 0.69 | 1.17 | Whether the shoot crosses the farm | 14314 |
| reg_f_firstmastitisUbroYellow | 1.04 | 0.84 | 1.51 | First line choice of mastitis treatment is streptomycin/ framycetin | 15162 |
| reg_f_muckspreader1 | 1.04 | 0.84 | 1.49 | Use of a contract muck spreader | 14473 |
| reg_f_manure_9yes | 1.04 | 0.74 | 1.8 | Use of other manure management | 14351 |
| reg_f_pneum_vacc1 | 1.04 | 0.85 | 1.45 | Routine use of calf respiratory vaccination | 14655 |
| reg_f_badger2 | 1.04 | 0.8 | 1.59 | Perceived badger population on the farm | 13939 |
| reg_f_manure_8yes | 1.03 | 0.79 | 1.61 | Use of separator for muck management | 14545 |
| reg_f_rook2 | 1.03 | 0.85 | 1.44 | Perceived rook population on farm | 15600 |
| reg_f_whichclinrespTetracycline | 1.03 | 0.83 | 1.49 | First line choice of pneumonia treatment is tetracycline | 14297 |
| reg_f_nsaidcalve1 | 0.97 | 0.71 | 1.16 | Routine use of anti-inflammatories at calving | 14669 |
| reg_f_manure_5yes | 0.97 | 0.69 | 1.19 | Muck heap in field used | 15226 |
| reg_f_hostfarmwalk1 | 1.03 | 0.84 | 1.44 | Whether a farm walk has taken place on the farm in the last 2 years | 14724 |
| reg_f_firstmastitisOrbeninLA | 0.97 | 0.52 | 1.44 | First line choice of mastitis treatment is cloxacillin | 16658 |
| reg_f_wean2 | 0.97 | 0.65 | 1.24 | Age of weaning | 12976 |
| reg_f_market1 | 1.03 | 0.84 | 1.43 | Whether animals are taken to market | 14828 |
| reg_f_calveclean.L | 1.03 | 0.86 | 1.36 | How often the calving pens are cleaned out | 14240 |
| reg_f_manure_6yes | 1.03 | 0.85 | 1.43 | Use of muck heap on concrete | 12981 |
| reg_f_abcalve1 | 1.03 | 0.8 | 1.54 | Routine use of antibiotics following a difficult calving | 15179 |
| reg_f_total_cattle | 1.03 | 0.88 | 1.29 | Total number of cattle on farm | 15234 |
| reg_f_bedmilk_1yes | 0.97 | 0.69 | 1.22 | Use of sand in milking cow bedding | 14603 |
| reg_f_feedlorry1 | 0.97 | 0.71 | 1.19 | Whether feed lorries crossed any cattle yards when delivering | 14536 |
| reg_f_herd_size | 1.03 | 0.88 | 1.3 | Total milking cattle on farm | 15734 |
| reg_f_leptovaccY | 0.97 | 0.7 | 1.22 | Routine use of leptospirosis vaccination | 14241 |
| reg_f_whichclinrespOtherDdontknow | 1.03 | 0.71 | 1.81 | First line choice of pneumonia treatment is unknown | 15240 |
| reg_f_bedmilk_6yes | 0.97 | 0.58 | 1.39 | Use of green bedding for milking cows | 14250 |
| reg_f_pigeon2 | 0.98 | 0.74 | 1.17 | Perceived pigeon population on farm | 15773 |
| reg_f_calving_group1 | 1.02 | 0.78 | 1.52 | Whether calves were born in a group pen | 14519 |
| reg_f_manure_2yes | 0.98 | 0.69 | 1.25 | Use of a weeping wall for slurry management | 15317 |
| reg_f_organic1 | 0.98 | 0.61 | 1.38 | Whether the farm is organic | 14649 |
| reg_f_um_spray1 | 0.98 | 0.69 | 1.26 | Routine use of umbilical treatment at birth | 14147 |
| reg_f_fox2 | 0.98 | 0.74 | 1.21 | Perceived fox population on farm | 15565 |
| reg_f_clox_dct1 | 0.98 | 0.71 | 1.23 | Use of cloxacillin dry cow therapy | 14750 |
| reg_f_othergrazeY | 0.98 | 0.73 | 1.21 | Use of grazing land away from the farm | 14461 |
| reg_f_whichclinrespMacrolide | 1.02 | 0.83 | 1.34 | First line choice of pneumonia treatment is Macrolide | 15840 |
| reg_f_fram_dct1 | 1.02 | 0.81 | 1.39 | Use of framycetin dry cow therapy | 13613 |
| reg_f_timesfirstmast | 1.02 | 0.89 | 1.23 | How often clinical mastitis cases were treated with intramammary tubes | 12810 |
| reg_f_scc | 0.98 | 0.84 | 1.11 | Somatic cell count | 15570 |
| reg_f_trough_clean.L | 1.02 | 0.86 | 1.25 | Daily cleaning of water troughs in calf housing | 14955 |
| reg_f_deer2 | 1.02 | 0.81 | 1.37 | Perceived amount of deer on farmland | 15528 |
| reg_f_nsaiddiarr1 | 1.01 | 0.82 | 1.34 | Routine use of anti-inflammatories in cases of calf diarrhoea | 15523 |
| reg_f_acf1 | 0.99 | 0.76 | 1.22 | Use of automatic cluster flushing | 16112 |
| reg_f_heifers_waste1 | 0.99 | 0.73 | 1.25 | Whether waste milk has been fed to heifers | 14992 |
| reg_f_manure_3yes | 0.99 | 0.74 | 1.24 | Use of a metal slurry store | 15682 |
| reg_f_daystreatmast | 1.01 | 0.87 | 1.21 | Number of days mastitis is treated for | 16019 |
| reg_f_rat2 | 1.01 | 0.82 | 1.3 | Perceived rat population on farm | 15792 |
| reg_f_nsaidmast | 1.01 | 0.88 | 1.2 | Routine use of anti-inflammatories in cases of mastitis | 15387 |
| reg_f_halocur1 | 0.99 | 0.77 | 1.24 | Routine use of treatment to prevent cryptosporidiosis | 15260 |
| reg_f_manure_1yes | 1.01 | 0.81 | 1.33 | Use of slurry lagoon | 14497 |
| reg_f_pheasant2 | 0.99 | 0.72 | 1.29 | Perceived pheasant population on farm | 14788 |
| reg_adult1 | 1.01 | 0.79 | 1.31 | Sample collected from the milking cow environment | 14004 |
| reg_f_drytrough.L | 1.01 | 0.85 | 1.22 | How often the dry cow water trough was cleaned out | 16126 |
| reg_f_bvdvaccY | 1.01 | 0.79 | 1.33 | Routine use of BVD vaccination | 13955 |
| reg_f_lungvaccY | 0.99 | 0.77 | 1.24 | Routine use of lungworm vaccination | 15196 |
| reg_f_foragetype_1yes | 1.01 | 0.72 | 1.44 | Feeding of clamp grass silage | 15213 |
| reg_f_diarrvaccY | 1.01 | 0.8 | 1.28 | Routine use of calf enteric disease vaccination | 15370 |
| reg_f_geographical_area3 | 1 | 0.76 | 1.3 | Geographical area 3 | 14587 |
| reg_f_premises2 | 1 | 0.79 | 1.29 | Number of farm premises | 15374 |
| reg_f_starling2 | 1 | 0.79 | 1.29 | Perceived amount of starling infestation | 15983 |
| main_fq | 1 | 0.67 | 1.42 | Total fluoroquinolone usage | 7315 |
| reg_f_milkgraze1 | 1 | 0.75 | 1.37 | Whether milking cows graze | 15098 |
| main_tet | 1 | 0.74 | 1.33 | Total tetracycline usage | 12200 |
| reg_f_bedmilk_4yes | 1 | 0.66 | 1.56 | Use of paper waste for milking cow bedding | 15625 |
| reg_f_foragetype_6yes | 1 | 0.7 | 1.46 | Feeding of straw | 15070 |
| reg_f_aiexternal1 | 1 | 0.79 | 1.28 | Artificial insemination performed by external company | 13663 |
| reg_f_bedmilk_3yes | 1 | 0.78 | 1.27 | Use of sawdust as milking cow bedding | 15820 |
| reg_f_outsource1 | 1 | 0.72 | 1.4 | Whether heifer calves are reared by a different enterprise | 15969 |

**Table S5:** Showing the variables identified as associated with resistance for each model and the effect on their coefficients when the model is re-run with sceptical priors.

**Amoxicillin**

| **Variable** | **Odds ratio** | **Lower CI** | **Upper CI** | **Odds ratio using sceptical priors** | **Lower CI using sceptical priors** | **Upper CI using sceptical priors** | **Description** |
| --- | --- | --- | --- | --- | --- | --- | --- |
| reg_s_housed_outdooroutdoor | 0.27 | 0.2 | 0.37 | 0.28 | 0.2 | 0.38 | Sample collected from pasture |
| reg_preweaned_heifer1 | 1.99 | 1.29 | 2.98 | 1.99 | 1.29 | 2.95 | Sample collected from the environment of a pre-weaned heifer |
| main_s_temp | 1.91 | 1.57 | 2.35 | 1.9 | 1.56 | 2.34 | Average monthly temperature |

**Ciprofloxacin**

| **Variable** | **Odds ratio** | **Lower CI** | **Upper CI** | **Odds ratio using sceptical priors** | **Lower CI using sceptical priors** | **Upper CI using sceptical priors** | **Description** |
| --- | --- | --- | --- | --- | --- | --- | --- |
| reg_preweaned_heifer1 | 4.13 | 2.79 | 6.46 | 4.12 | 2.79 | 6.41 | Sample collected from the environment of a pre-weaned heifer |
| main_s_temp | 2.14 | 1.63 | 2.87 | 2.12 | 1.61 | 2.82 | Average monthly temperature |

**Streptomycin**

| **Variable** | **Odds ratio** | **Lower CI** | **Upper CI** | **Odds ratio using sceptical priors** | **Lower CI using sceptical priors** | **Upper CI using sceptical priors** | **Description** |
| --- | --- | --- | --- | --- | --- | --- | --- |
| reg_preweaned_heifer1 | 1.95 | 1.46 | 2.51 | 1.95 | 1.46 | 2.51 | Sample collected from the environment of a pre-weaned heifer |
| main_s_temp | 1.53 | 1.32 | 1.77 | 1.52 | 1.32 | 1.77 | Average monthly temperature |

**Tetracycline**

| **Variable** | **Odds ratio** | **Lower CI** | **Upper CI** | **Odds ratio using sceptical priors** | **Lower CI using sceptical priors** | **Upper CI using sceptical priors** | **Description** |
| --- | --- | --- | --- | --- | --- | --- | --- |
| main_total_mg_pcu | 1.47 | 1.01 | 2.13 | 1.46 | 1 | 2.11 | Total antibiotic usage |
| main_strep | 0.76 | 0.6 | 0.97 | 0.76 | 0.6 | 0.97 | Total streptomycin usage |
| reg_s_rain | 1.26 | 1.08 | 1.47 | 1.26 | 1.08 | 1.46 | Average monthly rainfall |
| reg_s_housed_outdooroutdoor | 0.24 | 0.15 | 0.35 | 0.24 | 0.15 | 0.35 | Sample collected from pasture |
| reg_s_calving_now1 | 2.02 | 1.1 | 3.29 | 2.02 | 1.1 | 3.29 | Sample collected during the calving season |
| reg_preweaned_heifer1 | 3.39 | 2 | 5.82 | 3.41 | 2.01 | 5.8 | Sample collected from the environment of a pre-weaned heifer |
| main_s_temp | 1.98 | 1.55 | 2.55 | 1.97 | 1.55 | 2.52 | Average monthly temperature |

**Supplementary Figures**

**Figure S1**: Flow diagram to demonstrate the various stages used to arrive at the final dataset used in analysis of risk factors for sample-level *bla*_CTX-M_ *E. coli* positivity.

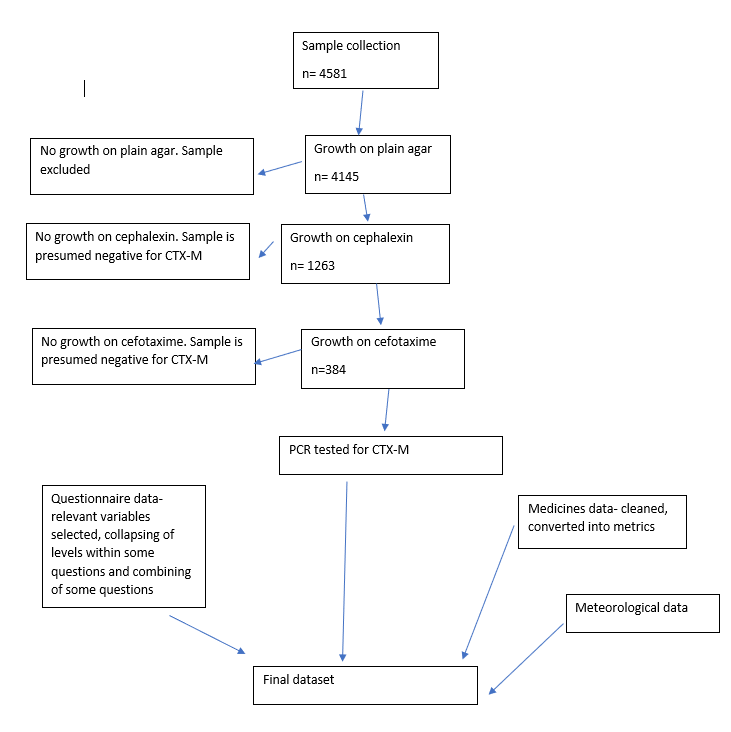

**Figure S2:** Effect of minimum *bla*_CTX-M_ *E. coli* prevalence (q) on the output of the risk factor model using the total dataset. Variable codes are: “preweaned_heifer”, Sample collected from a pre-weaned heifer (Calf sample); “s_temp”, Average monthly temperature; “weaned_heifer”, Sample collected from a weaned heifer (Heifer sample); “s_housed_outdoor”, Sample collected from pastureland; “f_foragetype_3”, Maize silage used on farm.

### A: The effect on the model estimates for different values of q.

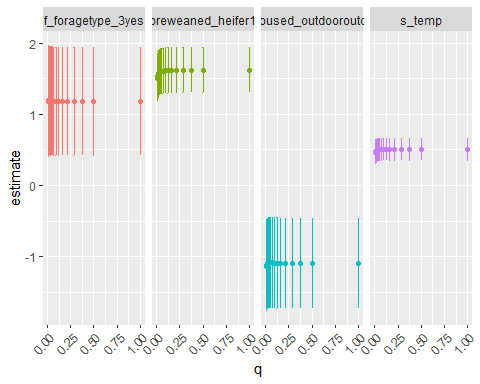

### B: Effect on the predictive accuracy of the model with different values of q, measured as the area under the Receiver Operating Characteristic Curve

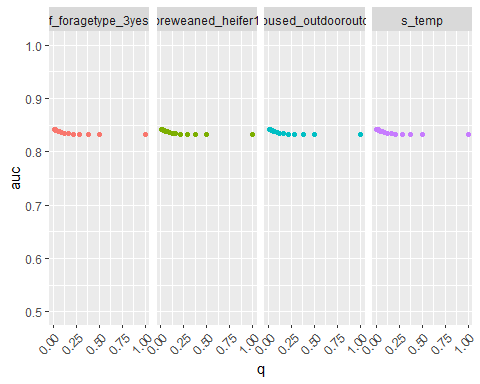

**Figure S3**: Initial questionnaire. This was completed by the researcher in the presence of the farmer at the time of consent and the first farm visit.

  
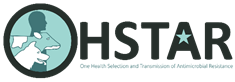

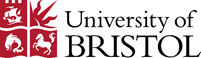

**The Risk Factors for Antibiotic Resistance on Dairy Farms Study**

**Initial questionnaire**

| **Farm name** |
| --- |
| **Address** |
| **Name of main contact** |
| **Main contact position** |

**Farm information**

| How many hectare (or acres) of land are on this holding? | hectares/acres |
| --- | --- |
| How much of this land is grazed? | hectares/acres |
| Are there any public footpaths, roads or other public access routes on your land? |  |
| Does the public have any right of access  through grazing land or to other areas where cattle are kept or walked? |  |
| Do the cattle kept here use any other grazing land?  If so, where , and does the public have access to it? |  |

**Do any of your cattle get grazed or wintered elsewhere?**

| **Name of site** |
| --- |
| **Postcode/location**    **Size of grazing land (if not included in above holding)** |

**How many people work on this farm?**

|  | **Brief description of role** |
| --- | --- |
| **Full-time** |  |
| **Part-time** |  |
| **Family members** |  |

**Are any other species kept on this farm or linked holdings?**

| **Species** | **Approx no of animals** | **In contact with dairy cattle?** | **Comments** |
| --- | --- | --- | --- |
| Sheep |  |  |  |
| Goats |  |  |  |
| Pigs |  |  |  |
| Poultry |  |  |  |
| Horses/donkeys |  |  |  |
| Dogs |  |  |  |
| Cats |  |  |  |

**Cattle demographics**

|  | **Number** | **Comments** |
| --- | --- | --- |
| Total cattle currently on site |  |  |
| Total dairy herd |  |  |
| Cows currently in milk |  |  |
| Preweaned dairy calves  (0-8wks) |  |  |
| Weaned dairy calves  (8wks-12m) |  |  |
| Dairy heifers 12-24 months |  |  |
| Dairy heifers over 24 months, not yet calved |  |  |
| Dairy cows |  |  |
| Total other cattle (give details) |  |  |

**How many animals were brought on to your land from elsewhere in the last year?** Include those who were temporary visitors such as bulls.

|  | **Private Market** | **Another Holding** | **Other, own herd** | **Other** |
| --- | --- | --- | --- | --- |
|  | **State number of animals and month(s) of arrival** | | | |
| **Dairy Calves**  Preweaned (under 8 weeks) |  |  |  |  |
| **Dairy Calves**  8 weeks to 12 months |  |  |  |  |
| **Dairy Heifers**  12-24 months |  |  |  |  |
| **Dairy Heifers**  Over 24 months, not yet calved |  |  |  |  |
| **Dairy Cows** |  |  |  |  |
| **Bulls** |  |  |  |  |
| **Beef cattle**  Give lifestages and month of arrival |  |  |  |  |
| **Other species in close contact with cattle (please give details)** |  |  |  |  |

**Fertility indices**

|  | **Average figures 12 month rolling** | **Comments** |
| --- | --- | --- |
| Average age at first calving | months |  |
| Calving to conception interval | days |  |
| Average dry period length | days |  |
| Do you use AI? |  |  |
| Do you use natural service? |  |  |
| Average number of serves/conception |  |  |
| 100 day in-calf rate |  |  |
| How frequently do you have a routine fertility visit? |  |  |

**Milk production**

|  | **Number** | **Comments** |
| --- | --- | --- |
| Total annual milk sales |  |  |
| Milk price average over 12 months | ppl |  |
| 305 day milk yield (12 month rolling average) | per cow |  |
| Somatic cell count (12 month rolling average) | per ml |  |
| Bactoscan (12 month rolling average) |  |  |
| Days in milk (12 month rolling average) |  |  |
| Number of milkings per day |  |  |
| Milking system used |  |  |
| Which company do you milk record with? |  |  |
| Percentage of cows in which antibiotic dry-cow tubes were used at dry-off in the last year | % |  |
| Percentage of cows in which systemic antibiotics were used at dry-off in the last year? | % |  |
| Clinical mastitis cases | per 100 cows/year |  |
| Percentage of clinical mastitis that occurs in the first 30 days of lactation | % |  |
| How is the milking routine adapted when high SCC cows are present? | | |

**Preventative healthcare**

| **Disease** | | **Routinely vaccinated for?** | **Routinely tested?** | **Herd status (+ve/-ve/**  **unknown)** | **Accredited as free from?** |
| --- | --- | --- | --- | --- | --- |
| Bovine Viral Diarrhoea | |  |  |  |  |
| Infectious Bovine Rhinotracheitis | |  |  |  |  |
| Leptospirosis | |  |  |  |  |
| Johnes | |  |  |  |  |
| Salmonella | |  |  |  |  |
| Huskvac/Lungworm | |  |  |  |  |
| Calf Respiratory Diseases (note vaccine used) | |  |  |  |  |
| Clostridial disease (Blackleg/7in 1) | |  |  |  |  |
| Pasteurella/Mannheimia | |  |  |  |  |
| Diarrhoea (eg Rotavec corona) | |  |  |  |  |
| Ringworm | |  |  |  |  |
|  | **Comments** | | | | |

**Lifespan of Dairy Cattle**

| Cull rate (excluding TB) | % | Comments |
| --- | --- | --- |
| Culling rate due to TB | % |  |
| Death rate | % |  |
| Average life span |  |  |

**Figure S4**: Final questionnaire. This was completed by the researcher during a telephone call with the farmer within two months of the last visit to the farm.

 
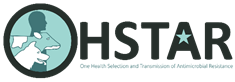

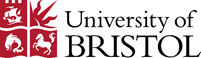

**One Health Selection and Transmission of Antimicrobial Resistance**

**Dairy farms: Final questionnaire**

|  |  | **Details** |
| --- | --- | --- |
| Do any of the people who work on your farm live or work on any other farms? | Yes/no |  |
| Do any of the people who live on your farm live or work on any other farms? | Yes/no |  |
| Staff turnover – how many new staff have started in the last 18 months?      How many have left?      Include temporary workers. | Started          Left |  |

**Herd size**

|  | **Number** | **Comments** |
| --- | --- | --- |
| Total cattle currently on site |  |  |
| Total dairy herd |  |  |
| Preweaned dairy heifer calves (calves still on milk) |  |  |
| Weaned dairy heifer calves (up to 12m) |  |  |
| Dairy heifers 12-24 months |  |  |
| Dairy heifers over 24 months, not yet calved |  |  |
| Other calves (preweaned) |  |  |
| Older beef cattle |  |  |
| Bulls |  |  |

**What other animals have been kept on your farm in the last 18 months?**

| **Species** | **Number of animals** | **If they were brought on farm in the last 18 months:** | | **Comments** |
| --- | --- | --- | --- | --- |
|  |  | **When did they arrive?** | **Where did they come from?**  Market  Another holding  Own holding  Abroad  Other |  |
| Sheep |  |  |  |  |
| Goats |  |  |  |  |
| Pigs |  |  |  |  |
| Poultry |  |  |  |  |
| Horses/donkeys |  |  |  |  |
| Dogs |  |  |  |  |
| Cats |  |  |  |  |

**How many cattle were brought on to your land from elsewhere in the last 18 months?**

Include those who were temporary visitors such as bulls.

| ***State number of animals and month(s) of arrival*** | **Private Market** | **Another Holding** | **Other, own herd** | **Other** |
| --- | --- | --- | --- | --- |
| Preweaned dairy calves (0-8wks) |  |  |  |  |
| Dairy heifers (8wks-12m) |  |  |  |  |
| Dairy heifers 12-24 months |  |  |  |  |
| Dairy heifers over 24 months, not yet calved |  |  |  |  |
| Dairy cows |  |  |  |  |
| Other calves |  |  |  |  |
| Older beef cattle |  |  |  |  |
| Bulls |  |  |  |  |

**Fertility**

| What is your calving pattern? | Tight block  Loose block  Most of the year but take a few months off  All year consistently  All year but more at certain times  Other |
| --- | --- |
| During which months did cows calve last year? |  |
| **Average figures 12 month rolling** | |
| Average number of serves/conception |  |
| Average age at first calving | months |
| Did the heifers in this study have AI? |  |
| Did the heifers in this study have natural service? |  |

**Milk**

|  | **Number** | **Comments** |
| --- | --- | --- |
| Total annual milk sales | Litres |  |
| Milk price average over 12 months | ppl |  |
| 305 day milk yield (12 month rolling average) | per cow |  |
| Somatic cell count (12 month rolling average) | per ml |  |
| Bactoscan (12 month rolling average) |  |  |
| Number of milkings per day |  |  |
| Milking system used (eg herringbone, abreast, rotary, robot) |  |  |
| Percentage of cows in which antibiotic dry-cow tubes were used at dry-off in the last year | % |  |
| Percentage of cows in which injectable antibiotics were used at dry-off in the last year? | % |  |

**Death**

| Cull rate (excluding TB) | % | Comments |
| --- | --- | --- |
| Culling rate due to TB | % |  |
| No. cow deaths on farm last 12 months |  |  |

**Figure S5**: Sensitivity of *bla*_CTX-M_ *E. coli* detection at different minimum prevalence of *bla*_CTX-M_-positive *E. coli* (q) as a proportion of total *E. coli* in a sample

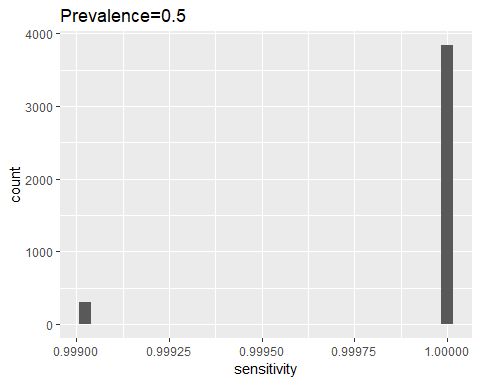

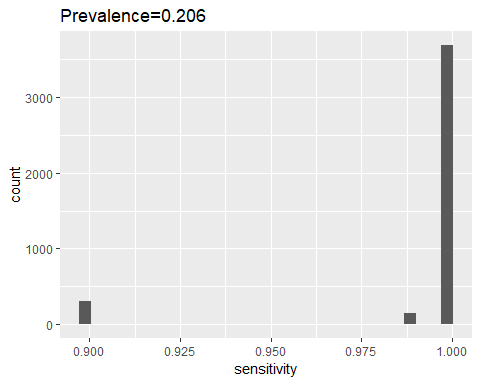

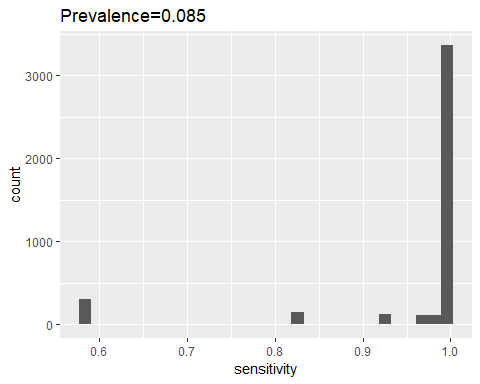

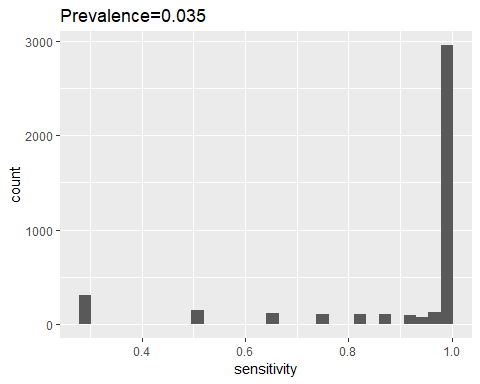

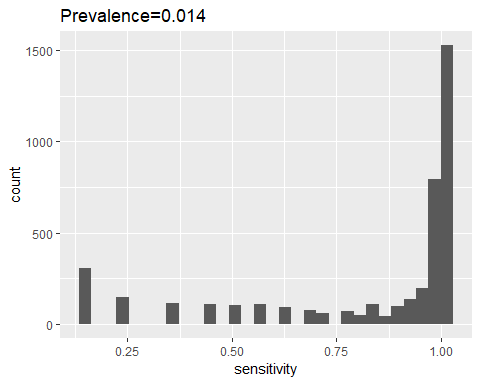

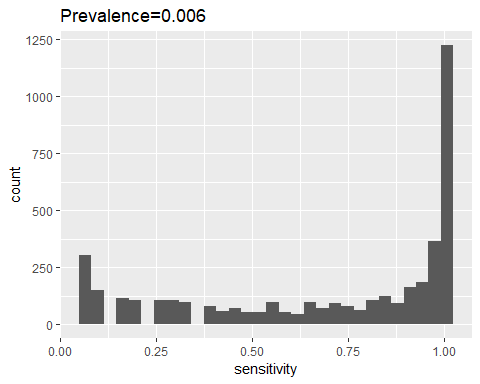

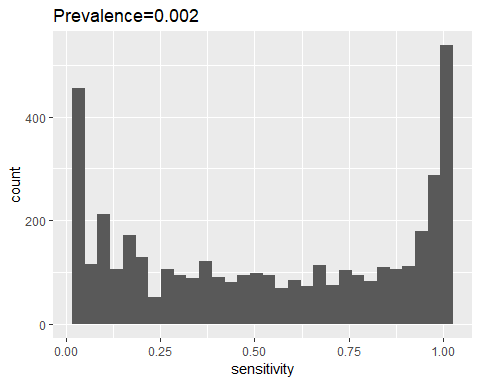

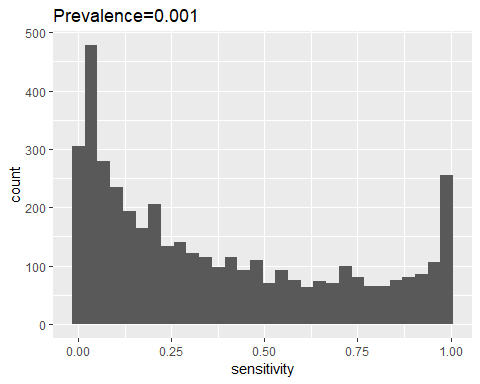

**Figure S6:** Trace plots from the Bayesian models of the variables that are identified as associated with resistance. All showing evidence of good convergence.

**Amoxicillin**

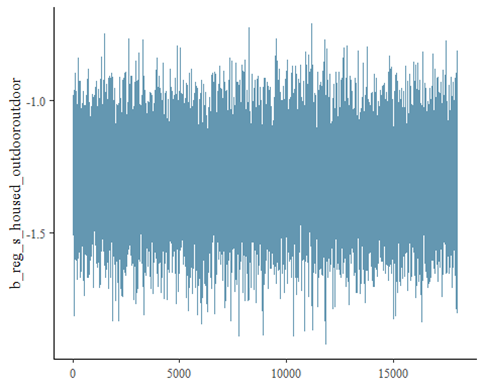

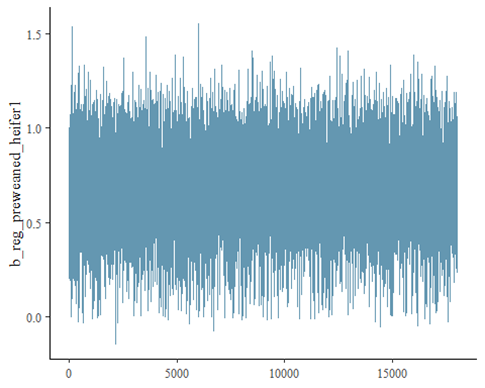

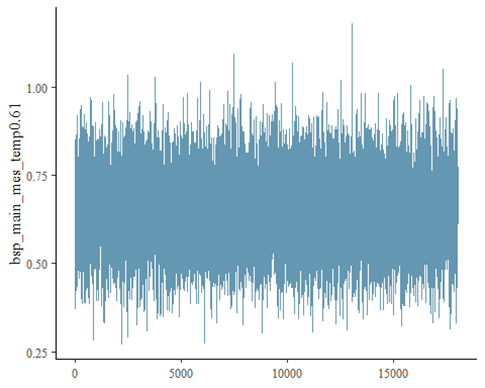

**Ciprofloxacin**

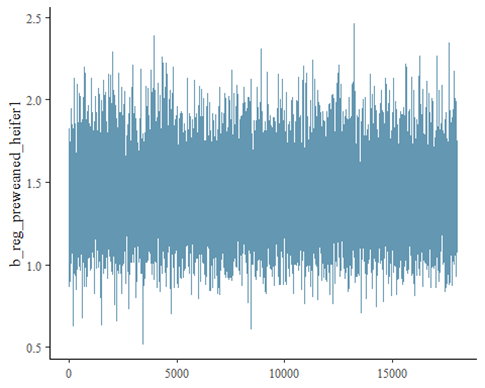

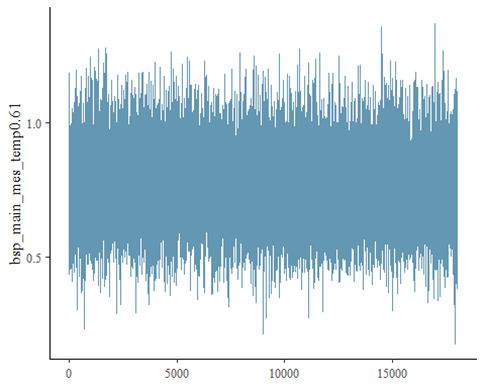

**Streptomycin**

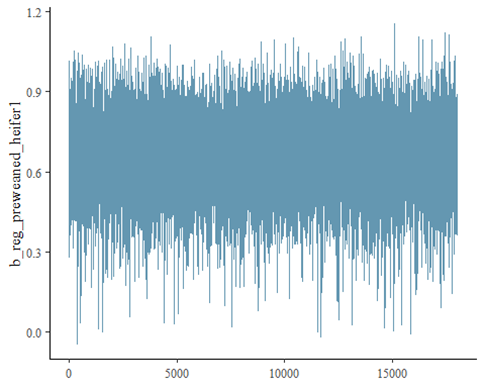

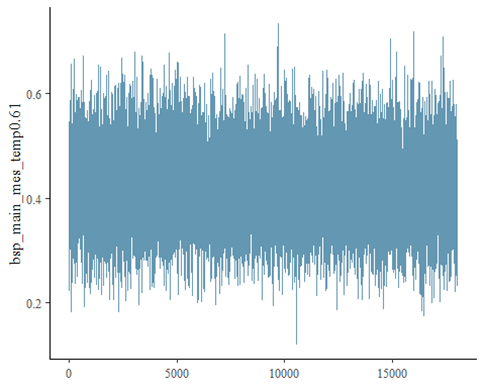

**Tetracycline**

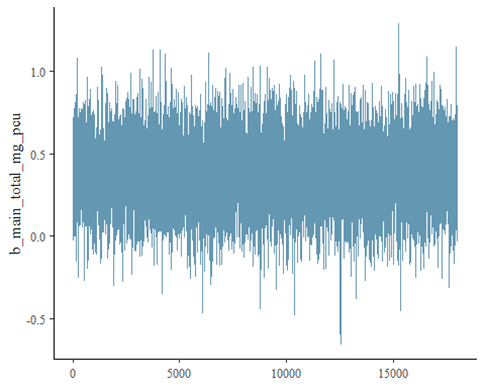

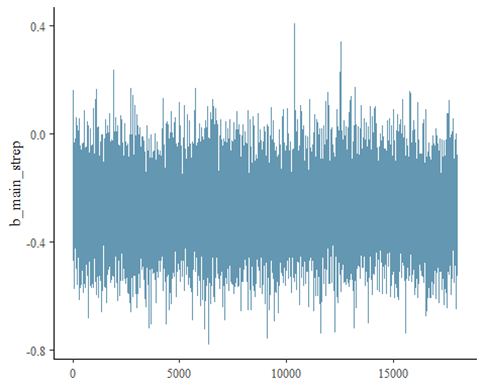

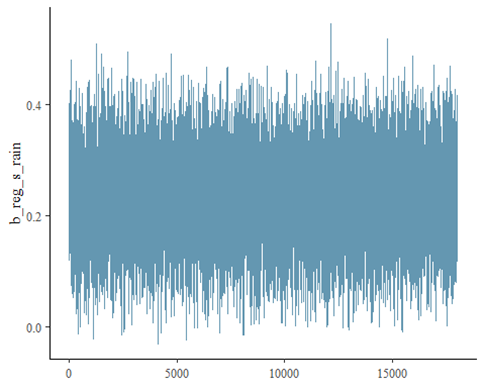

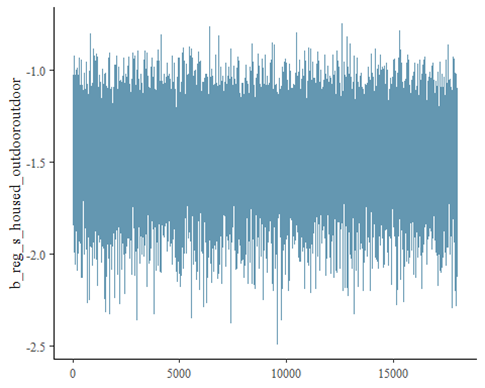

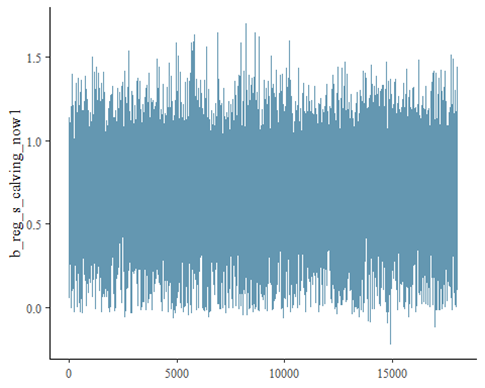

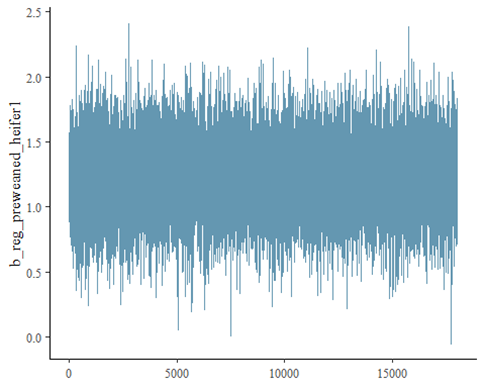

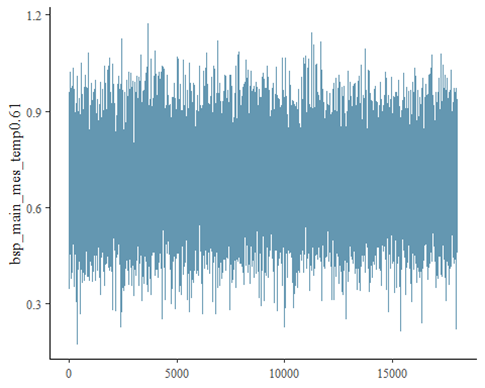

**Figure S7:** Posterior distributions with medians and 95% credible intervals of the variables that are associated with resistance.

**Amoxicillin**

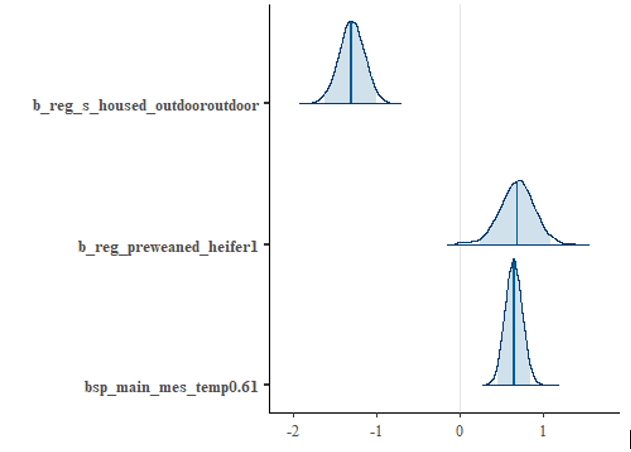

**Ciprofloxacin**

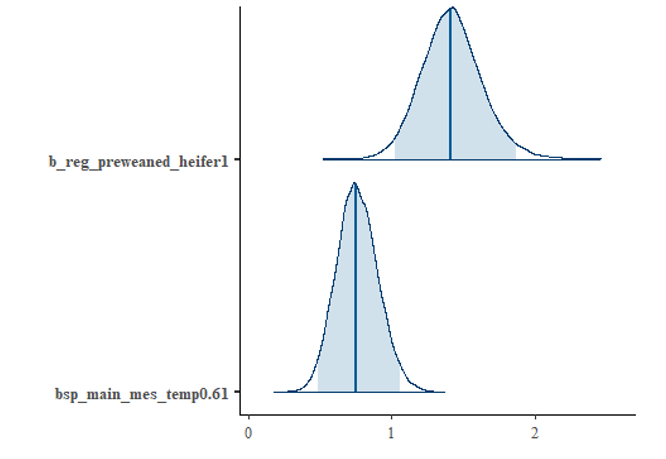

**Streptomycin**

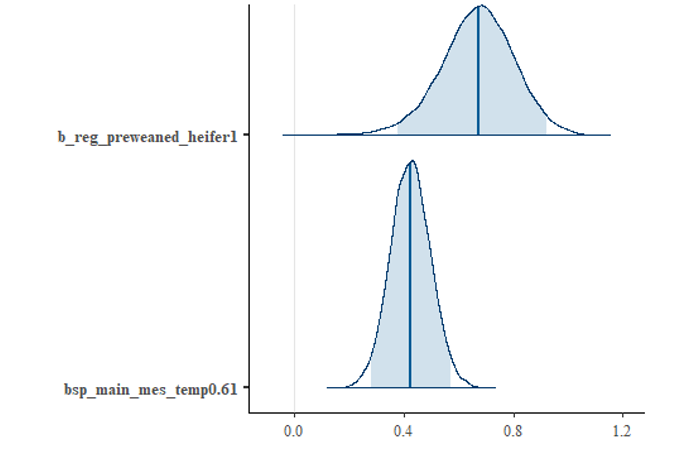

**Tetracycline**

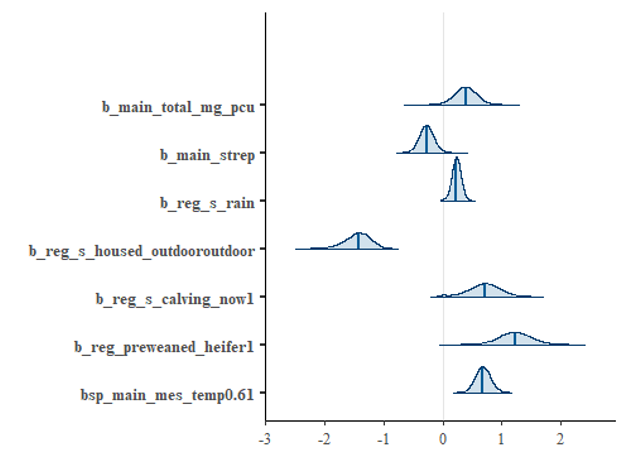
